# Redefining the connectome: A multi-modal, asymmetric, weighted, and signed description of anatomical connectivity

**DOI:** 10.1101/2022.12.19.519033

**Authors:** Jacob Tanner, Joshua Faskowitz, Andreia Sofia Teixeira, Caio Seguin, Ludovico Coletta, Alessandro Gozzi, Bratislav Mišić, Richard F. Betzel

## Abstract

The macroscale connectome is the network of physical, white-matter tracts between brain areas. The connections are generally weighted and their values interpreted as measures of communication efficacy. In most applications, weights are either assigned based on imaging features–e.g. diffusion parameters–or inferred using statistical models. In reality, the ground-truth weights are unknown, motivating the exploration of alternative edge weighting schemes. Here, we explore a multi-modal (combining diffusion and functional MRI data) regression-based, explanatory model that endows reconstructed fiber tracts with directed and signed weights. Benchmarking this method on Human Connectome Project data, we find that the model fits observed data well, outperforming a suite of null models. The estimated weights are subject-specific and highly reliable, even when fit using relatively few training samples. Next, we analyze the resulting network using graph-theoretic tools from network neuroscience, revealing bilaterally symmetric communities that span cerebral hemispheres. These communities exhibit a clear mapping onto known functional systems. We also study the shortest paths structure of this network, discovering that almost every edge participates in at least one shortest path. We also find evidence of robust asymmetries in edge weights, that the network reconfigures in response to naturalistic stimuli, and that estimated edge weights differ with age. In summary, we offer a simple framework for weighting connectome data, demonstrating both its ease of implementation while benchmarking its utility for typical connectome analyses, including graph theoretic modeling and brain-behavior associations.

## INTRODUCTION

The human connectome refers to the complete set of fiber tracts that link brain regions to one another [1]. It can be reconstructed non-invasively from diffusion weighted images using tractography algorithms [2, 3].

The connectome is of great interest to a number of scientific communities. Cognitive processes are supported by distributed, brain-wide networks [4, 5] and many neuropsychiatric disorders are thought to be disorders of dysconnectivity [6, 7]. Mapping connectomes and understanding their organizing and operational principles [3, 8–11] is also a key aim of network neuroscience [12] – the nascent discipline that focuses on modeling and analyzing brain data (micrographs, MR images, electrophysiological recordings) as networks.

The connectome is an example of an “anatomical” or “structural” network, in that the edges all represent physical, material pathways [13–16]. In anatomical networks, connections are usually associated with weights.

In human tractography data, these weights are frequently assigned based on diffusion parameters – e.g. fractional anisotropy, mean diffusivity, or streamline counts – and are interpreted as measures of white-matter fiber integrity.

A related but distinct approach involves mapping “effective connectivity” [17, 18]. This approach uses recordings of brain activity – usually functional MRI data – to model the directed influence exerted by one gray matter region on another. Effective connectivity can be estimated using measures of temporal precedence, such as Granger causality [19–21] or transfer entropy [22, 23], which assess whether errors/uncertainty in the predicted activity of one region can be improved by including information about the history of another. This class also includes dynamic causal models [24–30], which seek the underlying circuit that, given a generative model, can best explain observed brain activity.

These approaches for mapping anatomical and effective connections have pronounced limitations and trade off their utility with one another. For instance, diffusion-/tractography-based measures are incapable of resolving directionality – i.e. the weight of the tract from region *i* to *j* is identical to that of *j* to *i*. Also, these same measures are indirect assessments of myelination status (those note [31–34]) and not necessarily informative about synaptic efficacy or influence.

Measures of effective connectivity also have limitations. Precedence-based measures are typically estimated between all pairs of brain regions and not restricted to those for which a structural connection exists. This is a limitation shared by causal models, as well. In the case of causal models, the space of possible network parameters is huge, even for very small networks. This makes fitting causal models computationally costly [35]. Although recent approaches have helped reduce this burden, making it possible to fit cortex-wide networks, scaling beyond approximately 10^2^ nodes is uncommon. Even for these small networks, the computational cost remains demanding enough that investigating individual differences–i.e. fitting subject-specific models–is prohibitive [36, 37].

Here, we present a multi-modal, explanatory model for estimating the weights of structural connections. In the spirit of diffusion-/tractography-based models, ours preserves the brain’s sparse white-matter architecture. However, rather than assign structural weights based on diffusion/imaging parameters, we assign weights based on the parameters of multi-linear regression models. These models are fit independently for each region, *i*, and predict that region’s future activity based on the weighted histories of its connected neighbors. This allows us to fit asymmetric and signed edge weights for networks of hundreds of nodes in a matter of seconds.

Our manuscript aims to explore this model and its network properties, positioning it as an intermediate method, situated between tractography-based weighting schemes and computationally expensive inferential techniques. To this end, we find that the model predicts fMRI BOLD activity at a rate greater than chance even when using a relatively small amount of data (approximately 1% of samples). We show that the inferred edge weights exhibit subject specificity and are aligned, broadly, with known functional systems, despite the fact that edge weights exhibit imperfect alignments with interregional correlations (see Fig. S1). Further, we show that this network exhibits bilaterally symmetric, hemisphere-spanning communities and a shortest path structure that involves most edges (in contrast with streamline-weighted networks that use only 15% edges in its shortest paths backbone). Taking advantage of the directed nature of links, we find evidence of robust asymmetries in connection weights and regions’ connectivity profiles (incoming *versus* outgoing connections). Finally, in two applications we show that the inferred edge weights systematically reconfigure during movie-watching and across the human lifespan. Collectively, these observations suggest that this model is a practical alternative to existing edge-weighting schemes, and effectively endows anatomical connections with functional properties, thereby opening up avenues for future exploration and applications.

## FITTING AND BENCHMARKING ASYMMETRIC, WEIGHTED, AND SIGNED STRUCTURAL CONNECTIVITY

Brain regions are linked to one another *via* whitematter fiber tracts. The topology and edge weights of this network constrains interareal communication and shape patterns of spontaneous activity. Most studies set the weights of structural connections equal to microstructural properties estimated from diffusion weighted images and tractography, e.g. fractional anisotropy or streamline derivatives, or infer them using computationally demanding generative models. However, the ground truth connection weights are unknown, motivating the exploration and benchmarking of alternative weighting schemes.

Here, we use a model-based framework for assigning weights to existing structural connections. Briefly, we follow existing modeling work [38–41] and assume that at time *t* the state of region *i* (level of fMRI BOLD activity) is a function of its neighbors’ states at time *t* – 1 plus an offset (bias). That is:

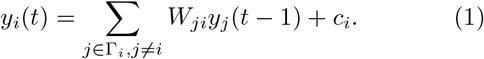

Where Γ_*i*_ is region *i*’s connected neighbors. We use linear regression and ordinary least squares to estimate the parameters *W_ji_* and *c_i_* separately for each node *i* (Fig. 1a + b). Thus, the resulting matrix 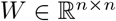, is sparse and preserves *exactly* the binary structure of white-matter connectivity (Fig. 1c). However, the weights, which are obtained from regression, can take on either positive or negative valence, whereas weights are typically positive only for connectomes inferred from dMRI and tractography. We note also that this network is directed–i.e. in general, *W_ij_* = *W_ji_*.

**FIG. 1.**
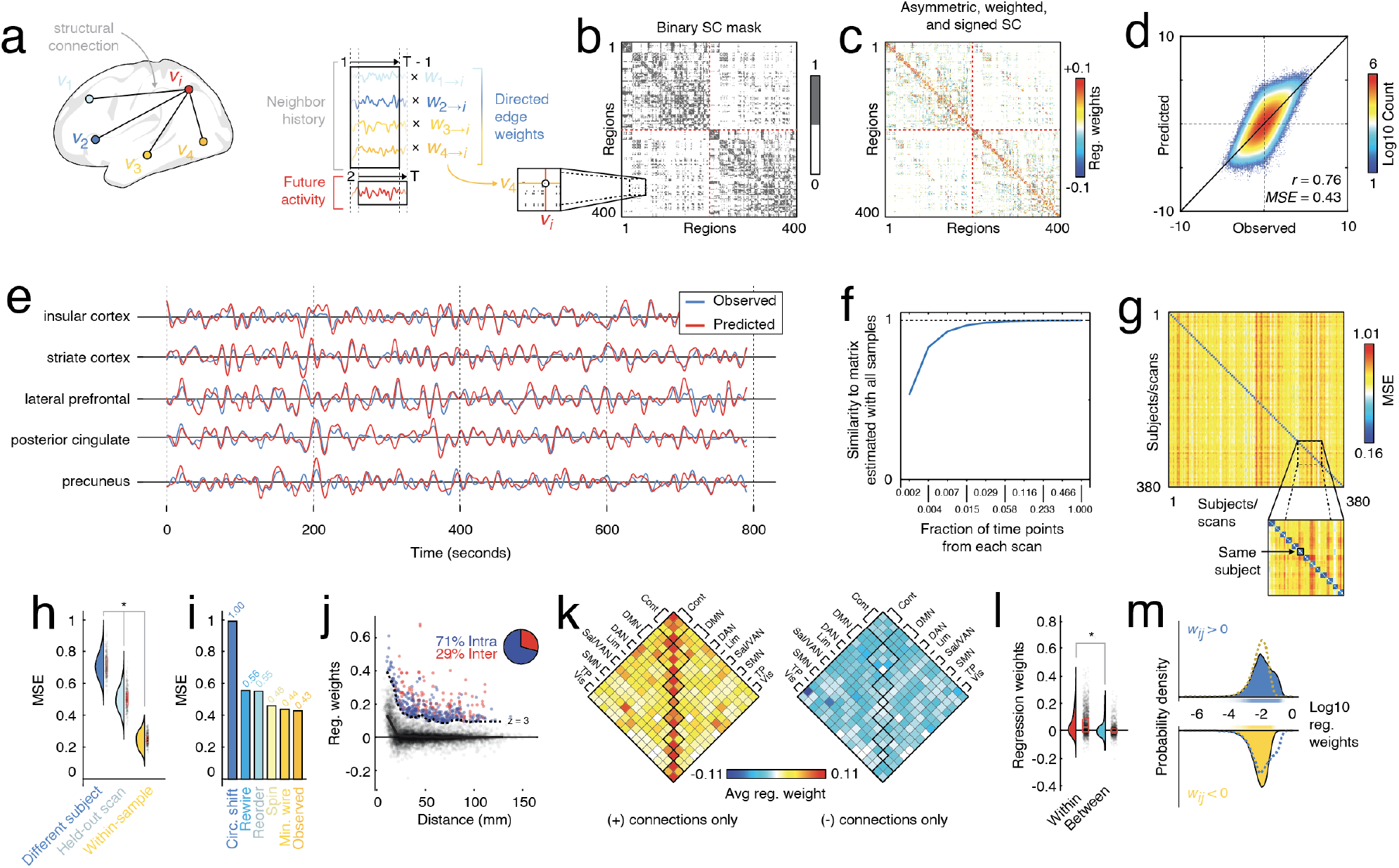
Fitting and characterizing weighted, signed, and asymmetric structural connectivity. (*a*) Here, we used linear model to estimate regression (edge) weights. For a given region *i*, we identified its structurally connected neighbors and used their past activity to predict node *i*’s future activity. This procedure results in a series of regression weights; one weight is associated with each neighbor. (*b*) Those weights were then entered into the binary structural connectivity “mask”. When this procedure was repeated for all *i* ∈ [1,…, 400] it generates a weighted, signed, and asymmetric matrix whose nonzero entries are masked by structural connections (see panel *c*). This matrix also has dynamic interpretations; a vector of brain-wide activity can be multiplied into the matrix to make a prediction of the activity at the subsequent time point. (*d*) Two-dimensional histogram showing observed and predicted activity, pooled across all participants and scans. The colors indicate number of samples in any bin. (*e*) Examples of observed and predicted activity for five select regions in a single subject and scan. (*f*) Similarity of regression weights (edge weights) as a function of amount of data. Note that units on *x*-axis are expressed as fraction of time points in scan, where the total number of frames was 1099. (*g*) Weights fit using scans from subject *s* are better at predicting activity in held-out scans from s than other subjects, *s*’. In this figure, the blocks along the diagonal are 4 × 4 and represent the four resting-state HCP scans. (*h*) Distributions of errors in model fit, grouped by whether the model is being used to predict activity in a scan of a different subject, the held-out scan from the same subject, or one of the scans used in fitting the weights. (*i*) Comparison of model errors using the observed network with a minimally wired network, one in which rows/columns were randomly reordered, and another in which time series were circularly shifted (independently across regions, scans, and subjects). (*j*) Relationship of regression weights and Euclidean distance. We also identified edges whose regression weights were much stronger than expected (those above the dashed line). (*k*) Distribution of regression weights across canonical brain systems. (*l*) Comparison of within- and between-system regression weights. (*m*): Comparison of positive and negative weights.

In this section, we report the results of the fitted model. That is, we describe basic features of the asymmetric, weighted, and signed connectome and contrast them with a connectome in which weights are defined using a commonly used metric–i.e. streamline density (streamline count divided by geometric mean of regional volumes) [3, 42].

We fit the model at the group level using pooled time series data from 95 participants from the Human Connectome Project [43] (HCP100UR, five subjects excluded due to incomplete data or quality issues) and a group-averaged binary SC matrix [44]. We found that, at the group level, the model performed well (correlation between observed and predicted activity from individual scans, *r* = 0.76 ± 0.03; mean squared error, *MSE* = 0.43 ± 0.05; Fig. 1d + e). We also found that the model weights stabilize with relatively few samples. Specifically, we randomly sub-sampled an equal number of frames from each participant and scan and used those frames to estimate the connection (regression) weights. We repeated this process 100 times while varying the number of samples from 2, 4, 8, 16, 32, 64, 128, 256, 512, to 1099. We found that even with approximately 6% of the total number of samples (64 samples per scan), the estimated weights achieved a correlation with the fullsample weights of *r* = 0.993; Fig. 1f).

Brain activity dynamics and its correlation structure are deeply individualized [45, 46]. A good model of brain activity, therefore, must also exhibit subject specificity. To assess whether model performance was, indeed, subject specific, we estimated weights using three of every subject’s four resting state scans, and used those weights to predict the activity of the held-out scan (as well as the activity of all other scans and subjects; Fig. 1g). We found that the error (mean squared error) was lower for the held-out scans than for the scans of any other subjects (two-sample *t*-test; *p* < 10^-15^; Fig. 1h). Note that here, and in all subsequent single-subject/-scan analyses, we fit edge weights using the same group-representative connectivity mask. This ensures that any differences between individuals are not driven by differences in the underlying anatomical connectivity, but driven jointly by differences in edge weights and resting brain dynamics.

Next, we assessed whether the observed results, namely the model error, was consistent with chance. Accordingly, we compared the observed model fitness against null distributions obtained under five distinct null models [47]: 1) a *minimally wired null model* in which only the shortest (least-costly) connections are preserved (while preserving an equal number of connections), 2) a *reordered null model* in which nodes’ orders were randomly permuted, 3) a *“spin” model* in which nodes’ orders were randomly permuted but geometry preserved, 4) a *topological null model* in which nodes’ degrees (number of connections and predictors) were preserved, but edge placement randomized, and 5) a *temporal* null model in which regional fMRI BOLD time series were circularly shifted independently for each region and scan (see **Materials and Methods** for details related to these null models). In general, we found that the error (MSE) was significantly lower using the intact data than in any of the null models (two-sample *t*-test, *p* < 10^-15^; Fig. 1i).

Finally, we examined some of the basic properties of the weights fit at the group level. We found that both positive and negative connection weights decay monotonically with distance (Fig. 1j). However, for any given distance bin there was a range of edge weight values. Examining the most extreme (z-scored weight of *z* > 3 relative to the other edges in the same bin), we find they are dominated by intra-hemispheric connections (≈ 71). Although fewer in number, the remaining 29% of connections still exceeds the baseline rate of inter-hemispheric connections (19.5%). Next, we tested how positive and negative connections were distributed with respect to canonical brain systems [48]. We found that, within systems, connections tended to be strong and positive whereas negatively-weighted connections showed no clear preference for falling either within or between systems. Indeed, when we examine the weights of individual connections, rather than system averages, we still find that within-system weights tend to be stronger and more positive compared to between-system weights (two-sample t-test, *p* < 10^-15^; Fig. 1l) and that, in general, the mean positive connection is greater than the absolute mean negative (two-sample t-test, *p* < 10^-15^; Fig. 1m).

In the supplementary material we perform several additional tests. These include assessing model performance at different lags (Fig. S2), assessing the relative contributions of long *versus* short connections (Fig. S3), comparing the estimated edge weights with other measures of functional and structural connectivity (Fig. S1), assessing regional fitness (Fig. S4), assessing the impact of global signal regression on results (Fig. S5), confirming that the distance dependence of edge weights is preserved when we use curvilinear fiber length rather than Euclidean distance (Fig. S6), fitting edge weights with regularization (Fig. S7), and comparing the relative performance of the asymmetric, weighted, and signed matrix *versus* the fiber density matrix in as structural constraints for dynamic, neural mass models (Fig. S8).

In summary, we show that this simple regression framework reliably estimates structural connection weights and requires relatively few observations to do so. The inferred weights are subject-specific and result in model fitness that exceeds chance. The strongest weights are positive and concentrated within putative brain systems. Collectively, these results set the stage for further explorations of the asymmetric, weighted, and signed network and the implications of the newly defined edge weights for network analyses.

## MODULAR ORGANIZATION OF THE ASYMMETRIC, WEIGHTED, AND SIGNED CONNECTOME

One of the hallmarks of biological neural networks is that they are organized into densely connected sub-networks called “modules” or “communities” [10]. Although there is a shared correspondence between anatomical modules–defined from streamline-derived structural connectivity–and functional modules, the alignment has, historically, been inexact [49, 50]. Here, we examine the modular structure of anatomical connectivity with the newly-derived asymmetric, weighted, and signed connectome and compare its organization with the modular structure derived from a connectome in which edges are weighted based on streamline density.

To detect communities we optimized a signed variant of the modularity quality function [51] using the Louvain algorithm [52]. The output of the algorithm is sensitive to initial conditions and was optimized 1000 times for each of the two weighting schemes. In both cases we fixed the structural resolution parameter to *γ* = 1. We aggregated and compared these results by computing coassignment matrices for each connectome, tallying the frequency with which node pairs were assigned to the same module across all 1000 repetitions (Fig. 2a,b). For the sake of visualization, we also calculated consensus communities for each matrix (Fig. 2c,d). We then calculated the difference between the two co-assignment matrices (Fig. 2e). We found that communities in the asymmetric, weighted, and signed matrix exhibited reduced laterality [53], tending to span the cerebral hemispheres whereas communities detected using the fiber density matrix tended to be more lateralized (t-test *p* < 10^-15^; Fig. 2g). We note that these observations were anticipated, given that fMRI BOLD activity was involved in the estimation of structural connection weights.

**FIG. 2.**
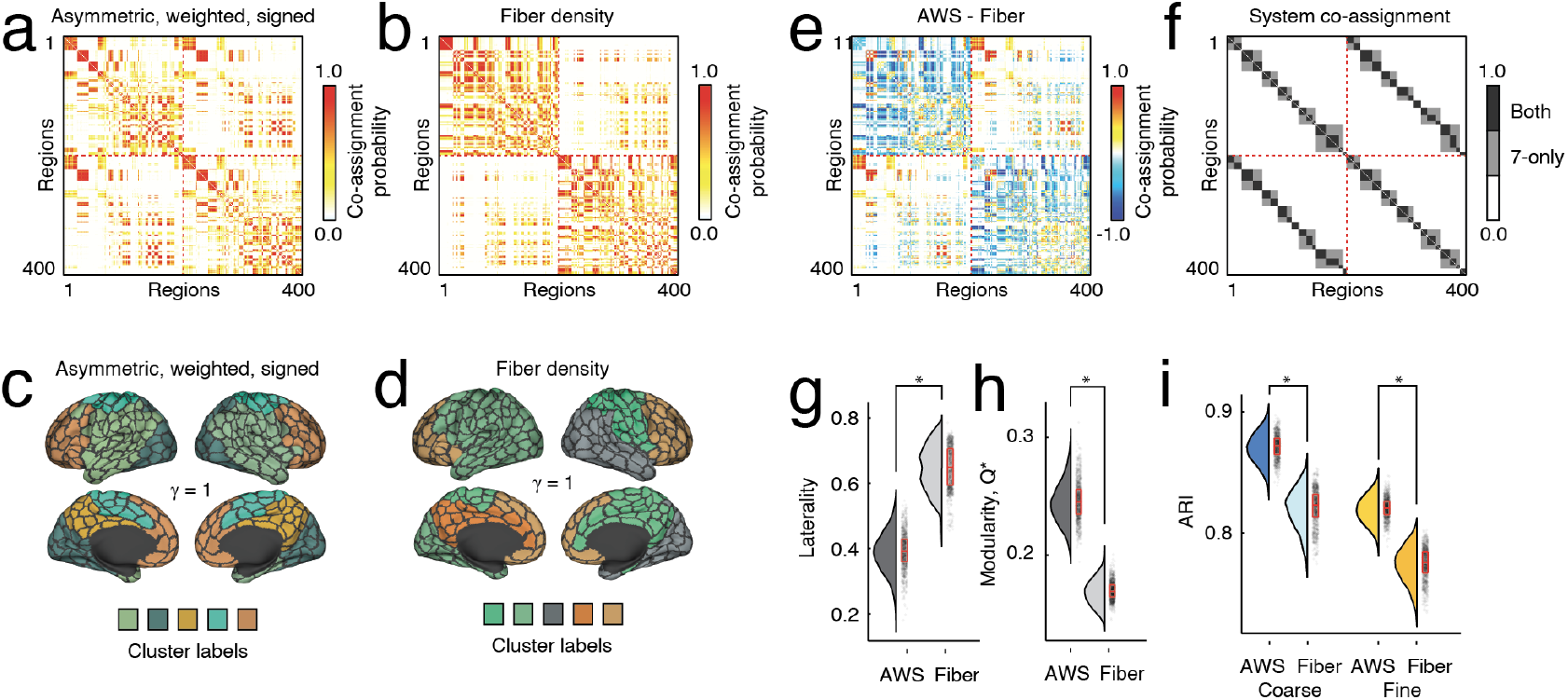
Comparing modular structure of structural networks. Modules are cohesive subnetworks – nodes that make more connections to other members of the same module than to other modules. Here, we compare the modular structure of network with original weights and the same network with weighted, directed, and signed edges. Here, we examine modules estimated with a fixed resolution parameter (*γ* = 1) but explore the multiscale modular structure in the supplement. Coassignment probability matrices for the inferred edge weight (*a*) and the fiber density matrices (*b*). (*c* and *d*) Consensus communities for both versions of weights. (*e*) Element-wise difference in module co-assignment. (*f*) System co-assignment matrix for reference. Black entries refer to pairs of brain regions that are assigned to the same system in both coarse and fine-scale system divisions. Gray entries are co-assigned to the same system for the coarse division, only. Comparison of laterality (*g*) and modularity (*h*) of detected modules. (*i*) Alignment of modules with respect to coarse and fine-scale system partitions. Each point in panels *g-i* represents a partition from one of 1000 runs of the Louvain algorithm for optimizing modularity.

Next, we asked whether the community structure of the asymmetric, weighted, and signed matrix was better aligned with functional connectivity (correlation structure of resting fMRI BOLD data) than the fiber density matrix and its system-level architecture. To address this question, we imposed canonical brain systems (coarse- and fine-scale intrinsic connectivity networks defined in [48]; Fig. 2f) on each matrix and calculated the induced modularity (*Q*\*). We found that the asymmetric matrix exhibited greater modularity than the fiber density matrix (*t*-test, *p* < 10^-15^; Fig. 2h). We also calculated the adjusted Rand index (ARI) between detected partitions and fine- and coarse-scale systems. ARI is a measure of partition similarity; larger values indicate that two partitions are more similar. For both the fine and coarse system partitions, we found that the ARI was greater when compared to partitions detected using the asymmetric and signed matrices than partitions detected using the fiber density matrices (two-sample *t*-tests; maximum *p* < 10^-15^; Fig. 2i).

For completeness, we also examined the multi-scale and hierarchical organization of communities, allowing for the resolution *γ* to vary [54] (Fig. S9). These results suggest that the asymmetric, signed, and weighted network exhibits community structure not limited to a single topological scale [55].

Notably, the results reported here were obtained using modularity maximization and a well-established null model [51]. We also explored an alternative “geographic” null model that has been used in network analysis of physical systems, e.g. granular materials [56–59] (details of this model are described in **Materials and Methods**; Fig. S10). Briefly, this null model preserves the binary network architecture exactly – the same presence/absence of links as in the observed network – but assigns a uniform weight across those edges. In general, we find that this null model generates results consistent with those described above, but also yields consensus communities that exhibit a striking correspondence to canonical brain systems (Fig. S11).

Additionally, we compared the detected modules to a recently aggregated set of “brain maps” [60] – annotations of brain regions that describe properties ranging from density of receptors to the relative expansion of brain areas across development and evolution. In general, we found evidence that the modules detected using the asymmetric, weighted, and signed network were more strongly enriched for these annotations compared to modules detected in the fiber density matrix (Fig. S12). This observation was true both at the level of the entire partition, but also at the level of individual modules.

Finally, we also repeated several of the analyses from this and the previous section using mouse anatomical connectivity data made available by the Allen Brain Institute [13] that was reprocessed and parcellated into *N* = 182 regions of interest (see **Materials and Methods** for processing details). Unlike dMRI and tractography, these connectivity data were acquired invasively using tract-tracing and are considered the “gold standard” for mapping of anatomical connectivity. The resulting networks have some shared features in common with networks reconstructed from dMRI [11], but are also directed and hyper-dense (of the possible 32942 connections, 32936 exist; 99.98% density). In parallel, we also analyzed fMRI data acquired from a cohort of *N* =18 anaesthetized mice, with data parcellated into the same *N* = 182 regions of interest (see **Materials and Methods** for details regarding acquisition, processing, and parcellation). In general, the results obtained using the mouse data were consistent with those reported using the human imaging data (see Fig. S13), including greater similarity between the community structure of the newly defined connectome weights and anatomically defined regions of mouse cortex.

## GRAPH THEORETIC PROPERTIES OF THE ASYMMETRIC, WEIGHTED, AND SIGNED CONNECTOME

In the previous section, we explored the modular architecture of the newly-derived asymmetric, weighted, and signed matrix, comparing it with analogous measures made on the fiber density matrix. Modular structure, however, is but one example of a network metric – it assesses a network’s organization at the “meso-scale”. However, other measures can be meaningfully applied to probe global (whole-network) and local (regional) properties. In this section, we investigate a subset of those measures.

First, we compared shortest-paths structure. Shortest paths in weighted networks refer to the least-costly route from a source node, *s*, to a target node, *t*. Typically, the length or cost of a shortest path is interpreted as a measure of communication capacity [61]; networks where the average shortest path is low (or the average reciprocal shortest path is large) are considered better-suited for communication.

To detect shortest paths we first mapped weights to costs. In the fiber density matrix, this mapping was accomplished by taking the reciprocal of an edge’s weight 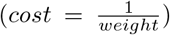 before applying a shortest paths algorithm. In the directed and signed network, however, we performed an additional step to rectify edge weights as the shortest paths algorithm is not compliant with negative edges. Briefly, we subtracted *min*(*β_ij_*) and added *ε* to every edge, where *min*(*β_ij_*) = −0.43 is the smallest (most negative) weight among all edges weights and *ε* = 0.0027 was the weight of the weakest edge in the fiber density network. This transformation ensures that all existing white-matter edges have weights that are nonzero and positive. Following this transformation, we used the reciprocal transform to map weights to cost.

The shortest paths matrices for both networks are shown in Fig. 3a,b. Strikingly, the number of steps in the least-costly paths was much greater for the fiber density matrix than for the asymmetric, weighted, and signed network (Fig. 3e,f). This likely is a consequence of the heavy-tailed fiber density distribution; because a small number of connections exhibit orders of magnitude stronger weights than the others, the cost of including those edges in shortest path is exceptionally small. From the perspective of the shortest paths algorithm, it is optimal to direct paths through these ultra low-cost edges, possibly even at the expense of direct connections [11, 62]. Further evidence for this claim comes from the shortest paths usage; in the fiber density matrix, the fraction of edges that are used in at least one shortest path is only 14.2% (Fig. 3c,d), whereas in the asymmetric, weighted, and signed networks, 97.5% of all edges get used at least once (Fig. 3).

**FIG. 3.**
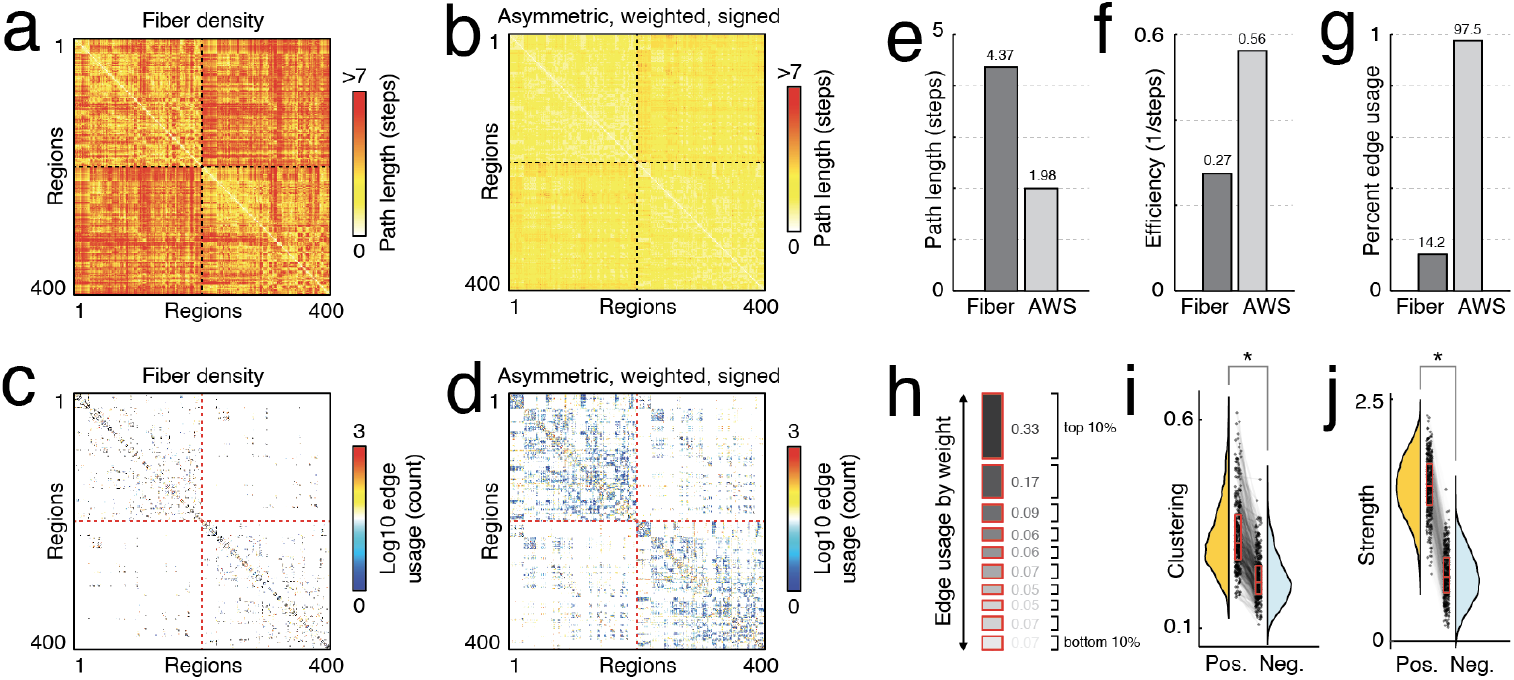
Network statistics for signed, weighted, and asymmetric matrix. (*a*) Path length (number of steps) between all pairs of nodes derived from the fiber density matrix 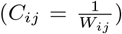. (*b*) Path length for new matrix 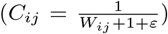. (*c*) Edge usage matrix for fiber density network. (*d*) Edge usage matrix for new matrix. Panels *e, f,* and *g* compare percentage of edges used in shortest paths, characteristic path length, and network efficiency between matrices. Panels *h* and *i* compare local clustering and strength (weighted degree) between the two matrices. We grouped edges into percentiles (deciles; lowest deciles include negative weights) based on their weights and calculated how frequently edges in each decile are involved in shortest paths. Panel *j* depicts the breakdown of edge usage by decile–e.g. top decile accounts for 33% of edges used on shortest paths.

The signed nature of the network means that we can also examine and compare properties of positive and negative edges to one another. That is, we can construct two versions of the same network: one in which nodes are linked *via* positive connections only and another with negative connections. Interestingly, we find that the positive network exhibits greater local clustering (paired sample *t*-test, *p* < 10^-15^; Fig. 3i). That is, positive connections tend to form dense triangles and cliques around nodes at a greater rate than negative connections. Additionally, we find that nodes’ positive weighted degrees (total weight of all incident positive connections) exceeds that of their negative strength (Fig. 3j).

In summary, we calculate a series of network statistics and show that their values differ, sometimes dramatically, depending on whether we weight edges using our regression-based framework or using more traditional diffusion/imaging parameters. In some specific cases, we find that statistics calculated on the asymmetric, weighted, and signed network are better aligned with our intuition about network function than statistics calculated based on fiber density. Collectively, these results underscore the impact of user decisions on network properties and our interpretation of network organization and function.

## ASYMMETRIES IN CONNECTION WEIGHTS

Due to technological limitations, structural connection weights estimated *in vivo* using diffusion imaging and tractography methods lack directionality–i.e. *W_ij_* = *W_ji_*. Here, however, the regression framework we use allows for asymmetries, such that the weights of incoming and outgoing connections can deviate from one another. In this section, we describe a select set of asymmetries in greater detail.

We measured asymmetry using a simple statistical test. Specifically, we identified pairs of regions whose weights were consistently asymmetric across 95 Human Connectome Project participants. This involved fitting weights independently for each subject and edge, and for every pair of nodes *i* and *j*, identifying connections where the distribution of asymmetry values, *W_ij_* – *W_ji_*, excluded zero (false discovery rate fixed at *q* = 0.05 resulting in *p_adj_* = 0.015; Fig. 4a,b) [63]. We found that, of 29024 possible connections, 8850 (approximately 30%) exhibited significant asymmetries.

**FIG. 4.**
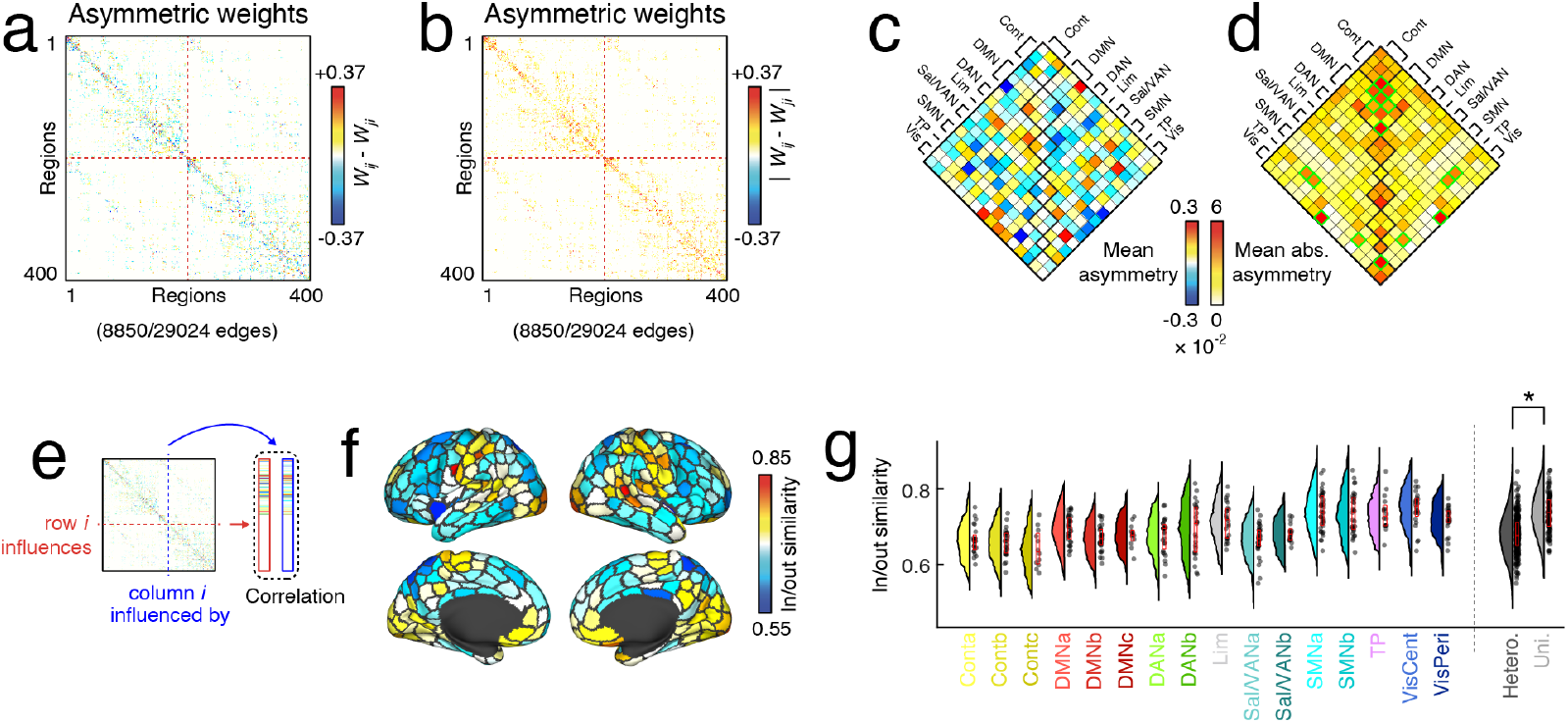
Asymmetries of influence between brain regions as assessed by inferred structural weights. (*a*) Significant asymmetries, and (*b*) absolute value of significant asymmetries were reorganized into system by system matrices (*c*) and (*d*) respectively. Note the increased absolute asymmetries within functionally defined systems in panel (*d*). As illustrated in the schematic in panel (*e*) next we measured in/out similarity as the correlation between incoming and outgoing weights per region. (*f*) Here we show an example of in/out similarity plotted to the brains surface. (*g*) Finally, we plot the per-system distribution of in/out similarity values across subjects. Boxplots on the right divide these systems into unimodal and heteromodal regions to show that there is more in/out similarity in unimodal systems.

Next, we asked whether edges whose weights were significantly asymmetric were preferentially concentrated within or between specific brain systems (Fig. 4c,d). To assess whether this was the case, we created a “mask” of edges that exhibited statistically significant asymmetries and aggregated (summed) these connections within and between every pair of systems. We performed this procedure first using the signed difference in edge weights and again using the absolute difference. These summed values were compared against a null distribution generated *via* a geometry-preserving null model [64]. We found that a number of system pairs exhibited greater than expected asymmetries, including connections that fall within control, default, and visual networks (ContC, DMNa, DMNc, central visual), as well as connections that fall between systems (ContC-DMNa, DMNa-DMNb, temporo-parietal and both ContC and DMNa, central visual and SMNb, and peripheral visual with DANb).

As a second measure of asymmetry, we compared the weights of nodes’ incoming and outgoing connection profiles – the extent to which its activity is predicted by *versus* predicts the activity of its neighbors. To do this, we calculated the linear product-moment correlation between vectors associated with row and column i in the asymmetric, weighted, and signed connectivity matrix (Fig. 4e). This procedure resulted in a single similarity score (correlation) for each brain region. In general, we found that in-out similarity was region-specific and varied between putative brain systems (Fig. 4f), with regions in sensorimotor systems exhibiting greater in/out similarity (Fig. 4g). Indeed, when we grouped systems based on unimodal (visual + somatomotor) and hetero-modal (all other systems) labels, we found that unimodal systems exhibited greater similarity (two-sample *t*-test, *p* < 10^-15^; Fig. 4h).

As a final test, we also identified node pairs, *i* and *j*, where sign(*W_ij_*) ≠ sign(*W_ji_*) (Fig. S14). We performed this analysis at the level of individual subjects and calculated the proportion of edges with an asymmetry of sign that fall either within or between brain systems (Fig. S14a). We repeated this analysis for every individual and found that between-system edges were more likely to exhibit an asymmetry of sign than within-system edges (two-sample *t*-test, *p* < 10^-15^; Fig. S14b).

Collectively, these results suggest that local asymmetries are well circumscribed by canonically defined brain systems. Additionally, our results suggest that asymmetry in regional incoming and outgoing connection weights run along a unimodal-heteromodal axis.

## WEIGHTS ARE MODULATED BY STATE

In the previous sections we validated and characterized structural networks whose edge weights were fit using a regression-based procedure. In those sections, all weights were fit to maximize the correspondence of predicted and observed activity during the resting-state. Here, we assess whether weights estimated at rest are dissociable from those estimated during movie-watching.

To address this question, we used resting-state and movie-watching data from the Human Connectome Project’s 7T dataset, focusing on a subset of 117 participants whose data passed quality checks and for whom all four scans were available. Using the same SC binary mask, we fit edge weights at the group level, pooling data across all subjects and scans to generate two asymmetric, weighted, and signed matrices: one based on resting-state and the other based on movie-watching data (Fig. 5a,b). In parallel, we also fit models at the level of individual subjects, pooling scans from the same subject to generate estimates of resting-state and moviewatching edges weights. In both cases, we found comparable performance (mean squared error) between both rest and movie data, with movies exhibiting slightly better performance than resting state (paired sample *t*-test; subject-level, *p* < 10^-15^; group-level *p* = 1.2 × 10^-12^; Fig. 5f-h).

**FIG. 5.**
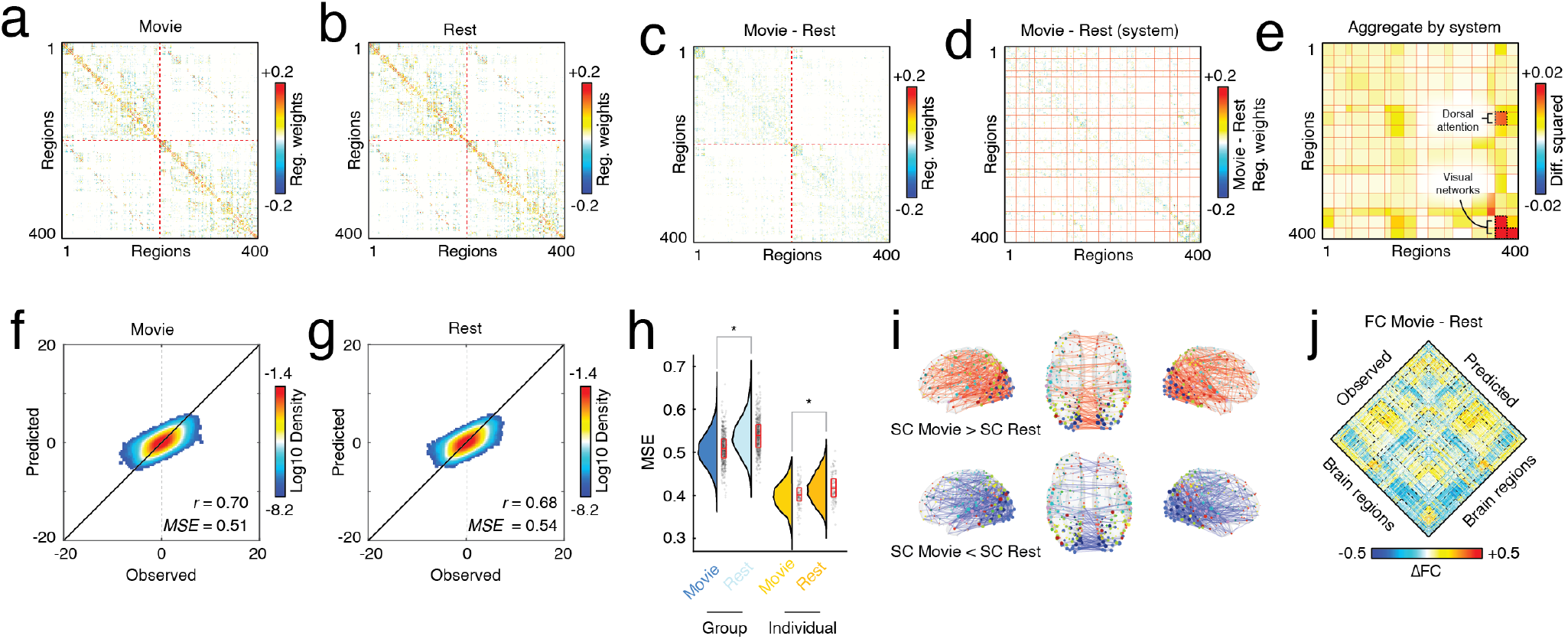
Comparing matrices fit to resting-state and movie-watching data. We analyzed 7T data from 117 participants in the Human Connectome Project. Using the same binary mask as in the previous sections, we fit edge weights for both conditions at the group level (pooling time series data across all subjects/scans) and individual level (pooling data from the same subjects). Weights for (*a*) group-level movie-watching state matrix and (*b*) group-level resting-state matrix. Panels *c* and *d* show the difference in edge weights (movie minus rest); rows and columns in panel c are ordered identically to panels *a* and *b*, whereas in panel *d* rows and columns are reordered by brain systems. (*e*) The average weight across existing connections between every pair of systems. Here, weights were squared prior to averaging. Panels *f* and *g* depict two-dimensional histograms of observed and predicted activity. (*h*) Model fitness when fit to pooled, group-level data (*left*) and individual data (*right*). (*i*) Edges whose difference between movie and rest are significantly greater or less than zero plotted in anatomical space. (*j*) Differences in functional connectivity (FC) between movie and rest for both the observed data (*left*) and the predicted (*right*).

First, we calculated the difference between edge weights for each subject and averaged the differences across subjects (Fig. 5c,d). At each edge, we performed a paired-sample *t*-test on the differenced edge weight distributions. We found that, out of *m* = 29204 total edges, 2463 exhibited significant state-dependent differences (multiple comparisons controlled for by fixing false discovery rate at *q* = 0.01 and adjusting the critical *p*-value, *p_adj_* = 8.5 × 10^-4^). Although these edges were distributed across the entire brain, they were significantly concentrated within a small subset of systems (Fig. 5e; dashed black borders around system blocks). Specifically, we found significant system-level effects within central and peripheral visual networks, from edges in the central visual network to the peripheral visual network (but not *vice versa,* and from the dorsal attention network (DANa) to the central visual network (spin test, false discovery rate fixed at *q* = 0.01, *p_adj_* = 1.6 × 10^-4^).

Projected into anatomical space, we find that, as expected, the connections that differ from rest to moviewatching tend to involve regions in visual networks (Fig. 5i). Interestingly, there are approximately as many connections whose weights increase from rest to movies as there are those that decrease, an effect that holds both within the visual networks (241 increases *versus* 218 decreases) but also across the entire brain (1292 increases *versus* 1171 decreases).

## DIFFERENCES IN THE WEIGHTED, SIGNED, AND DIRECTED CONNECTOME ACROSS THE HUMAN LIFESPAN

To this point, we have estimated the weights of asymmetric, weighted, and signed structural connections, described properties of the resulting network, exposed asymmetries in connections’ weights, and demonstrated that the weights are systematically modulated by task (rest *versus* movie). In this section, we investigate individual differences in connections’ weights and associate them with differences across the human lifespan (7-85 years).

To do so, we used data from the Nathan Kline Institute’s enhanced Rockland Sample [65], which included both diffusion weighted and functional MRI data for *N* = 542 participants. In-scanner head movement is known to vary systematically with age. To address motion-related concerns, we adopted the same conservative procedure as reported in Esfahlani *et al.* [66] for motion censoring. Specifically, for each of the remaining subjects, we dropped frames in which motion exceeded a pre-defined threshold (*FD* > 0.15 mm). We also dropped time points that were within two frames of any supra-threshold frame or failed for form a contiguous sequence of five frames or more. Following this procedure, we excluded any participant for whom the fraction of retained frames was fewer than 50% of their total number of frames. Collectively, these procedures left *N* = 474 participants with high-quality (low-motion) data for further analysis.

For each subject, we used the regression-based framework to fit weights to every structural connection, generating subject-specific asymmetric, weighted, and signed matrices. Note that, here, we restricted every subject to have the same binary set of consensus edges estimated from subjects aged 18-35 years; only edges’ weights varied across individuals. Prior to calculating age-related differences, we regressed out of each edge the following variables: sex (binary variable), gray matter volume, mean framewise displacement of all “low-motion” frames, and number of frames dropped due to motion contamination. Finally, using the residuals from this procedure, we calculated their linear product-moment correlation with subjects’ biological ages resulting in a (sparse) matrix of age-related correlations (Fig. 6a). For the sake of visualization, we show subsets of edges that pass uncorrected statistical tests (Fig. 6g-i).

**FIG. 6.**
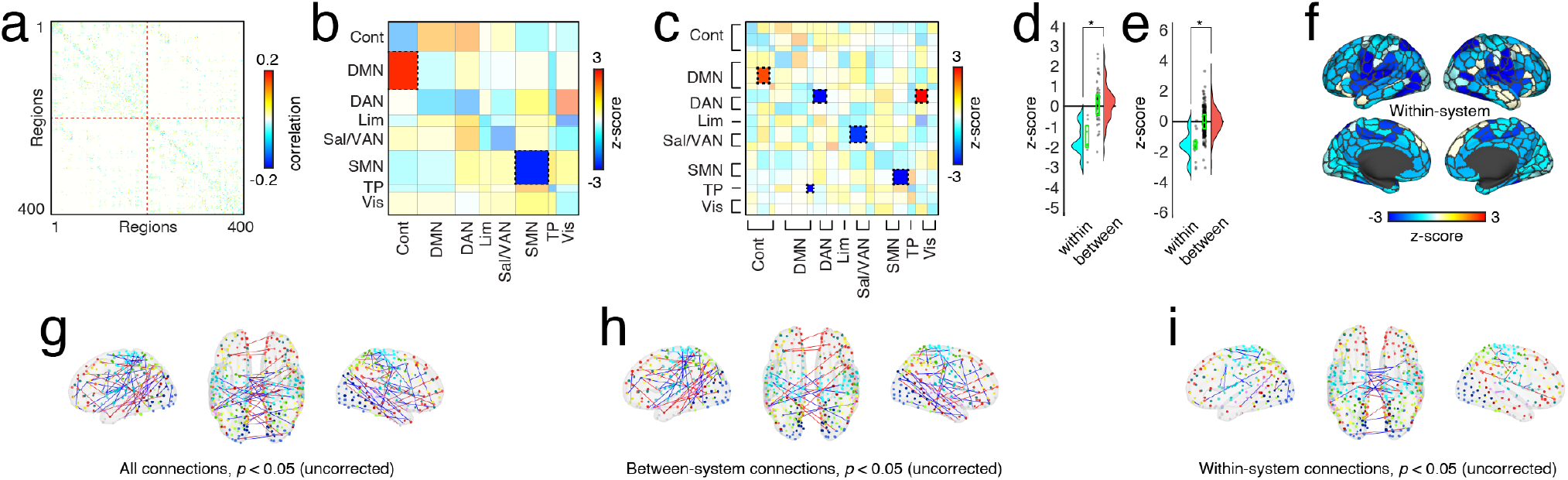
Age-related differences in inferred structural weights are more common within functional brain systems. (*a*) Edge level correlations with age. (*b* and *c*) Matrices of significant results for the coarse- and fine-scale system spin tests, respectively.(*d* and *e*) Boxplots comparing the z-scores from the spin-test null model with the real values. One boxplot displays within system z-score values, and the other displays between system z-score values. (*f*) Within-system z-scores from the fine-scale system spin test mapped onto cortical parcels. Panels *g-i* show individual edges that exhibit age-related correlations (*p* < 0.05; uncorrected). Blue and red edges correspond to edges whose weights decrease or increase with age, respectively. Panels *g-i* are presented for the sake of visualization only.

Previous studies have found that age-related differences in functional connectivity respect putative system boundaries [67–69]. Accordingly, we performed statistical tests at the level of systems [48]. Specifically, we calculated the mean correlation of all edges that fell between/within every pair of systems. We then compared these observed values with null distributions generated using “spin tests” (1000 repetitions). System pairs for whom the observed correlation exceeded that of the null were considered statistically significant (false discovery rate fixed at *q* = 0.05; *p_adj_* = 0.0088). In line with previous work, we found that age-related decreases in connection weight tended to concentrate within brain systems, whereas between-system weights were, generally, centered around a value of zero (two-sample *t*-test, *p* = 7.65 × 10^-13^; Fig. 6b-f). More specifically, at the coarse level, we found that connections within the somatomotor network significantly decreased their weight with age while connections from the default mode to the control network increased (Fig. 6b). The finer scale allowed us to better localize those effects while also discover new age-related differences. In particular, we found significant increases in connection weight from default mode B to control B, as well as an increase in connection weights from dorsal attention network A to the central visual module – an effect that had been previously obscured at the coarse scale. We also detected significant decreases in connection weight with age, concentrated within somatomotor network B, as well as previously undetected decreases within dorsal attention network A, salience/ventral attention network A, and from the temporo-parietal network to default mode C.

Altogether, these findings recapitulate well-known age effects that had been previously reported using functional connectivity data. However, our approach grounds these effects in anatomical connectivity, forming a multi-modal bridge between studies of anatomical and functional age-related differences and opening up avenues for future applied studies.

## DISCUSSION

The “correct” weighting of structural connections is not known. Most strategies for assigning edge weights do so based on microstructural or tractographic parameters. Though commonly used, the values of these parameters are, in general, misaligned with the interpretation that weights reflect the efficacy of inter-regional communication. Alternatively, a number of groups have proposed using measures of “effective” or “directed” connectivity, in which edge weights correspond to the magnitude of directed influence in a (biophysical) generative model of brain activity. However, due to computational complexity this approach is limited to applications involving small networks. Additionally, although fitting effective connections can be restricted to only those edges for which white-matter tracts were traced, most models are generally unconstrained and can place effective connections between disconnected brain regions. In short, there exists a plurality of methods for determining the weights of structural connections and the dominant methods each have specific strengths and weaknesses.

Here, we explored a simple regression-based model for endowing reconstructed fiber tracts with directionality and a signed weight. Benchmarking this method on Human Connectome Project data, we found that the model fit observed data well, outperforming a suite of null models. The estimated weights were highly reliable even when fit using relatively few training samples and exhibited marked subject specificity. We next analyzed the resulting network using tools from network neuroscience. These analyses revealed communities that spanned cerebral hemispheres and mapped clearly onto known functional systems. Almost every edge in this network was involved in at least one shortest path. We also found evidence of asymmetric weights, network reconfiguration during naturalistic movie watching, and age-related differences. We note that unlike biophysical and dynamic causal models, our weighting scheme is not generative– i.e. it cannot be used to generate new synthetic data. It is, however, explanatory and represents a means of weighting fiber tracts that is distinct from those most frequently used in network neuroscience. Collectively, the proposed framework presents opportunities for multiple follow-up studies and applications in other neuroscience disciplines.

### A new weighting scheme yields unique insights into brain network function

Over the past two decades, network neuroscience has led to a number of discoveries about the organization of brain networks. These include small-worlds [8], hubs [3] and rich clubs [9], modules and communities [10], and cost-efficient wiring [70]. These “canonical” findings have been observed not only in human brain networks reconstructed at the macroscale, but have been reported across phylogeny and at all spatial scales [71].

Despite this preponderance of converging evidence, the specifics of these findings often depend critically on whether or not to weight edges and the precise measure used. Consider the prototypical “small world” model of the brain, where long-distance edges are thought to represent shortcuts that allow signals to propagate long distances in relatively few hops. Although both weighted and binary networks exhibit small-world properties– relatively short path length and strong local clustering– when edges are weighted based on streamline density, the strong distance-dependence of streamlines ensures that virtually no long-distance connections are included in the network’s shortest path structure [11].

The example cited above represents just one instance where a processing decision (whether and how to weight structural connections) leads to different interpretations of brain network function, *vis a vis* role of long-distance connections in interareal communication. Interestingly, we make a similar observation here; when edges are weighted with regression coefficients and their signs rectified, we recover a shortest path structure in which most edges contribute to at least one shortest path and modules that are now better aligned with functional brain systems[72]. Our work even presents challenges for how we define connectomes [73]. When we refer to the connectome, we often imagine an enumerable and finite set of neural elements and connections [1, 74]. Here, however, edge weights are context dependent, impermanent, and vary with task/cognitive state. Essentially, the structural edges inherit features usually reserved for functional connections, placing our approach slightly at odds with the perspective that structural connections are fixed over short timescales (duration of typical scan session).

More importantly, the secrets of the connectome are far from unlocked. Existing data and methods have not unambiguously mapped structure to function, for example whether graph-theoretic measures can unveil functional properties of a brain (and which properties, specifically) is not clearly elucidated, and, although many studies have identified statistical associations between network properties and clinical, cognitive, behavioral, and developmental markers, the mechanisms that underlie those associations are largely unknown. Collectively, this motivates further neuroscientific exploration, both of new data/connectivity modalities [75] and experimental paradigms, as well as methodological frameworks.

### A network bridge between anatomical and functional connectivity

The edge-weighting scheme that we explore here can be used to help understand one of the central questions of network neuroscience: how does the brain’s anatomical connectivity constrain its function? Past studies have established a link between structural and functional connectivity [49]. Empirical findings have shown that insults to structural connections cause acute loss or reorganization of functional connectivity [76]. Even in intact brains, structural connection weights are correlated with their functional analogs and pairs of brain regions that are connected directly or *via* few processing steps have proportionally stronger FC [3]. In parallel, *in silico* dynamical systems models have used anatomical connectivity to constrain simulated brain activity [77, 78], generating synthetic fMRI data whose correlation structure can be compared with empirical FC or analytic estimates of interregional communication capacity [61, 79].

Here, rather than compare SC to FC, we incorporate functional information directly into the estimates of edge weights [80]. This process generates a singular network object whose fitness (a metric that, itself, can be interpreted as a measure of structure-function coupling) can be estimated globally as the total error between observed and predicted activity [81] or parsed into local (regional) error terms, analogous to recent approaches for linking anatomical and functional connectivity weights [66, 82]. We note however, that this approach is also distinct from most studies that report structure-function correspondence, in that we seek a set of parameters that maximizes that correspondence, whereas most studies report a correlation between functional data (FC or activity) and structural networks and their derivatives [83].

Here, we explore this alternative approach for tracking changes in structure-function correspondence in two contexts: comparing edge weights between rest and moviewatching conditions and identifying differences in edge weight across the human lifespan.

We find robust reconfiguration of edge weights during movie-watching. This observation is not new–a seemingly limitless number of studies have reported task- or state-induced changes in connection weights [84–87]. In these studies, however, it is the functional connections whose weights change, making it difficult to assess which structural connection facilitate those changes. Our approach addresses this issue directly; changes in regression weights at each structural edge allow for the formulation of targeted hypotheses about the roles of specific fiber tracts in a given task.

### Possible advantages of an asymmetric, weighted, and signed connectome

Throughout the study, we compare properties of the asymmetric, weighted, and signed network with a network in which edge weights represent fiber densities – a more traditional measure of structural connectivity. In general (and unsurprisingly), the properties of these networks are often dissimilar. That is, how we choose to weight a network’s edges can change its graph-theoretic profile and impact how we might interpret its function. Although we remain agnostic as to which weighting scheme is “superior”, we note that the asymmetric network both outperforms the fiber density network on a number of applications, and has properties that are better aligned with intuition.

For instance, we find that the modular structure of the asymmetric network tends to be less lateralized– i.e. modules are more likely to contain nodes from both hemispheres–than the fiber density network. This observation suggests that the reweighting of the network helps circumvent one of the peculiarities (or limitations) of community-detection methods applied to structural brain networks. Namely, because fiber densities and weights derived from tract-tracing experiments tend to be heavy-tailed and distance-dependent [11, 42] and because long-distance interhemispheric tracts are notoriously challenging to reconstruct from diffusion imaging data [88, 89], communities tend to be spatially contiguous and exhibit poor correspondence with systems/communities derived from functional recordings [49].

Additionally, we take advantage of recent advances in neuroinformatics to compare communities with brain maps–i.e. the regional or vertex-wide expression of genetic, transcriptomic, evolutionary, and developmental markers. We find that communities obtained from the asymmetric network tend to be significantly enriched for many of these markers to an extent above and beyond the communities obtained from the fiber density matrix. These observations suggest that the multi-modal network generated by endowing structural connections with functionally-relevant information tightens the link between network organization and brain-based markers.

### Interpreting signed edges in macroscale connectome

One of the unique features of the networks we construct here is that, unlike connectomes whose weights are determined based on microstructural/diffusion/tractography parameters, we obtain weights that are signed (can take on positive or negative values). How do we interpret this feature? What are possible underlying neurophysiogical mechanisms that explain the emergence of signed edges in large-scale networks?

At its core, the edge weights estimated here are statistical constructs; a negative weight from node *i* to *j* indicates that when the activity of *i* increases, the activity of *j* tends to decrease proportionally at the next time point. This statistical interpretation is in line with other frameworks for estimating effective connections [90]. Although it is tempting to ascribe “excitatory” and “inhibitory” labels to positive and negative edge weights, this terminology is typically reserved for cell-to-cell projections and, more specifically, pyramidal cells and interneurons, respectively. In general, the neurochemical (e.g. glutamate/excitatory and GABA/inhibitory) contributions to the diffusion MRI and fMRI BOLD signal are not easily parsed.

However, we can speculate about possible underlying mechanisms that support signed edges in large-scale networks. One possible explanation involves feed-forward or feedback inhibition [91], whereby excitatory interregional connections cause inhibition in their target region either by directly exciting local inhibitory interneurons, or by exciting local interneurons indirectly through connecting pyramidal cells. Indeed, recent studies have suggested that the balance of glutamate/GABA underlies the antagonistic (anti-correlated) activity of large-scale brain systems [92].

Irrespective of the underlying cause, the signed edges in the networks constructed here exhibit nonrandom organization in terms of their distribution across canonical brain systems and relationship to other network/geometric measures–e.g. clustering coefficient and fiber length. These features, of course, have implications for traditional network analyses and may require new methodologies in some cases. For instance, interregional communication models rely on shortest paths and diffusion dynamics to estimate communication efficacy [61]. However, these dynamics are not well-defined for networks with signed edges. Here, we circumvent this issue by offsetting edge weights, forcing negative connections to have small (but positive) values. However, there are likely many alternative strategies that embrace the signed and asymmetric nature of edges that could be explored in future studies [93].

### Limitations

This study has a number of limitations. Most notably, the primary results rely on the use of diffusion weighted MRI and tractography for reconstructing white-matter tracts. Although these methods are still used widely, they have well-documented drawbacks and biases that call into questions the verisimilitude of structural connectivity networks [94, 95]. We partially mitigate these concerns by replicating our findings using tract-tracing data made available through the Allen Brain Institute [13]. Unlike tractography, in which anatomical connections are inferred non-invasively, antero-/retrograde tract-tracing is considered the gold standard mapping large-scale connectivity [96]. We expect that advances in imaging, tractography algorithms [97, 98], and better alignment of multi-scale datasets will narrow the gap between tract-tracing and tractography in future studies [99].

Another possible limitation of the current study is its focus on neocortex only. Doing so necessarily ignores contributions from subcortical and cerebellar regions when modeling node-level time series. Adding additional regions is not computationally prohibitive and in principle could be addressed easily. However, new connections mean including additional explanatory terms in each regional multi-linear model and will, in general, lead to new estimates of edges’ weights. That is, the regression coefficients will vary with additional observations (new data) or new nodes. Future studies should investigate this explicitly.

### Future directions

Our study presents a number of opportunities for future studies. Among the most obvious and tantalizing opportunities is the empirical validation of edge weights estimated here. Datasets in which stimulation is paired with brain-wide recordings make it feasible to estimate directed influence between brain stimulus-target pairs of regions [100, 101]. These estimates could be compared directly to the connection weights inferred here. Relatedly, modeling studies have reported whole-brain effective connectivity [36] and asymmetries in communication patterns [63]. Here, we focused on characterizing properties of the network generated using our regression-based framework and contrasting it with a network of edges weighted based on fiber density; careful comparisons of this technique to extant, already-established methods should be carried out in the future.

Our preliminary findings using movie-watching data suggest that our weighting scheme may be suitable for detecting state-specific changes in structural connection weights. Future studies should explore the sensitivity of this approach for other state-based comparisons. We note that, unlike unthresholded functional connectivity, which, when used in a brain-wide association or casecontrol study results in *N*(*N* – 1)/2 comparisons, our approach results in much fewer connections [102]. Statistical tests only need to be performed for existing structural connections, possibly increasing the statistical power of these types of studies [103]. Additionally, while the edges we weight are structurally-defined and reflect best estimates of white-matter topology, we endow them with functionally relevant edge weights (derived from fMRI BOLD data). The multi-modal nature of this edge-weighting scheme may, in actuality, situate our approach somewhere between functional and anatomical connectivity. That is, it achieves what resting-state FC sometimes is assumed to be; namely, a functionally informative measure of anatomically connectivity [104].

Brain-wide association studies are familiar analyses in network neuroscience. In these studies, inter-individual variation in demographic, phenotypic, clinical, or behavioral data is associated with neural elements – usually node- or edge-level properties. In general, these types of studies are underpowered to detect small effect [102, 103]. This issue is especially problematic when tests are performed at the level of edges – for a network of N nodes, this means performing *N*(*N* – 1)/2 tests – which requires stringent corrections to control for multiple comparisons. Our approach may partly circumvent this issue (or at least reduce its severity); because anatomical connectivity is, in general, sparse, even if a statistical test is carried out at every edge, the total number of tests is far smaller than the upper limit. With fewer tests, the correction for multiple comparisons is less severe, potentially making it possible to resolve smaller effects.

## MATERIALS AND METHODS

In this section, we describe all four datasets that we analyzed. Briefly, they include three human MRI datasets: two from the Human Connectome Project and another from the Nathan Kline Institute. In addition, we also analyzed tract-tracing and functional MRI data from mice.

### Datasets: Human Connectome Project 3T resting-state and diffusion weighted MRI

The Human Connectome Project (HCP) 3T dataset [43] consists of structural magnetic resonance imaging (T1w), functional magnetic resonance imaging (fMRI), and diffusion magnetic resonance imaging (dMRI) young adult subjects, some of which are twins. Here we use a subset of the available subjects. These subjects were selected as they comprise the “100 Unrelated Subjects” released by the Connectome Coordination Facility. After excluding data based on completeness and quality control (4 exclusions based on excessive framewise displacement during scanning; 1 exclusion due to software failure), the final subset included 95 subjects (56% female, mean age = 29.29 ± 3.66, age range = 22-36). The study was approved by the Washington University Institutional Review Board and informed consent was obtained from all subjects.

A comprehensive description of the imaging parameters and image prepocessing can be found in [105]. Images were collected on a 3T Siemens Connectome Skyra with a 32-channel head coil. Subjects underwent two T1-weighted structural scans, which were averaged for each subject (TR = 2400 ms, TE = 2.14 ms, flip angle = 8°, 0.7 mm isotropic voxel resolution). Subjects underwent four resting state fMRI scans over a two-day span. The fMRI data was acquired with a gradient-echo planar imaging sequence (TR = 720 ms, TE = 33.1 ms, flip angle = 52°, 2 mm isotropic voxel resolution, multiband factor = 8). Each resting state run duration was 14:33 min, with eyes open and instructions to fixate on a cross. Subjects underwent 14 task fMRI scans over a two-day span. The fMRI data was collected with the same sequence parameters as the resting state fMRI. The fMRI runs consisted of working memory (5:01 min, 405 frames), gambling (3:12, 253), motor (3:34, 284), language (3:57, 316), social cognition (3:27, 274), relational processing (2:56, 232), and emotional processing (2:16, 176) tasks. Finally, subjects underwent two diffusion MRI scans, which were acquired with a spin-echo planar imaging sequence (TR = 5520 ms, TE = 89.5 ms, flip angle = 78°, 1.25 mm isotropic voxel resolution, b-vales = 1000, 2000, 3000 s/mm^2^, 90 diffusion weighed volumes for each shell, 18 b = 0 volumes). These two scans were taken with opposite phase encoding directions and averaged.

Structural, functional, and diffusion images were minimally preprocessed according to the description provided in [105], as implemented and shared by the Connectome Coordination Facility. Briefly, T1w images were aligned to MNI space before undergoing FreeSurfer’s (version 5.3) cortical reconstruction workflow, as part of the HCP Pipeline’s PreFreeSurfer, FreeSurfer, and PostFreeSurfer steps. Functional images were corrected for gradient distortion, susceptibility distortion, and motion, and then aligned to the corresponding T1w with one spline interpolation step. This volume was further corrected for intensity bias and normalized to a mean of 10000. This volume was then projected to the 2mm *32k_fs_LR* mesh, excluding outliers, and aligned to a common space using a multi-modal surface registration [106]. The resultant cifti file for each HCP subject used in this study followed the file naming pattern: `*_Atlas_MSMAll_hp2000_clean.dtseries.nii`. These steps are performed as part of the HCP Pipeline’s fMRIVolume and fMRISurface steps. Each minimally preprocessed fMRI was linearly detrended, band-pass filtered (0.008-0.008 Hz), confound regressed and standardized using Nilearn’s `signal.clean` function, which removes confounds orthogonally to the temporal filters. The confound regression strategy included six motion estimates, mean signal from a white matter, cerebrospinal fluid, and whole brain mask, derivatives of these previous nine regressors, and squares of these 18 terms. Spike regressors were not applied. Following these preprocessing operations, the mean signal was taken at each time frame for each node, as defined by the Schaefer 200 parcellation [48] in *32k_fs_LR* space. Diffusion images were normalized to the mean b0 image, corrected for EPI, eddy current, and gradient non-linearity distortions, and motion, and aligned to subject anatomical space using a boundary-based registration as part of the HCP pipeline’s Diffusion Preprocessing step. In addition to HCP’s minimal preprocessing, diffusion images were corrected for intensity non-uniformity with `N4BiasFieldCorrection` [107]. The Dipy toolbox (version 1.1) [108] was used to fit a multi-shell multi-tissue constrained spherical deconvolution [109] to the data with a spherical harmonics order of 8, using tissue maps estimated with FSL’s `fast` [110]. Tractography was performed using Dipy’s `Local Tracking` module [108]. Multiple instances of probabilistic tractography were run per subject [111], varying the step size and maximum turning angle of the algorithm. Tractography was run at step sizes of 0.25 mm, 0.4 mm, 0.5 mm, 0.6 mm, and 0.75 mm with the maximum turning angle set to 20°. Additionally, tractography was run at maximum turning angles of 10°, 16°, 24°, and 30° with the step size set to 0.5 mm. For each instance of tractography, streamlines were randomly seeded three times within each voxel of a white matter mask, retained if longer than 10 mm and with valid endpoints, following Dipy’s implementation of anatomically constrained tractography [112], and errant streamlines were filtered based on the cluster confidence index [113]. For each tractography instance, streamline count between regions-of-interest were normalized by dividing the count between regions by the geometric average volume of the regions. Since tractography was run nine times per subject, edge values were collapsed across runs. To do this, the weighted mean was taken with weights based on the proportion of total streamlines at that edge. This operation biases edge weights towards larger values, which reflect tractography instances better parameterized to estimate the geometry of each connection.

### Datasets: Human Connectome Project 7T resting-state and movie-watching data

The Human Connectome Project (HCP) 7T dataset [43] consists of structural magnetic resonance imaging (T1w), resting state functional magnetic resonance imaging (rsfMRI) data, movie watching functional magnetic resonance imaging (mwfMRI) from 184 adult subjects. These subjects are a subset of a larger cohort of approximately 1200 subjects additionally scanned at 3T. Subjects’ 7T fMRI data were collected during four scan sessions over the course of two or three days at the Center for Magnetic Resonance Research at the University of Minnesota. Subjects’ 3T T1w data collected at Washington University in St. Louis. The study was approved by the Washington University Institutional Review Board and informed consent was obtained from all subjects.

We analyzed MRI data collected from *N_s_* = 129 subjects (77 female, 52 male), after excluding subjects with poor quality data. Our exclusion criteria was as follows: where each spike is defined as relative framewise displacement of at least 0.25 mm, we excluded subjects who fulfill at least 1 of the following criteria: greater than 15% of time points spike, average framewise displacement greater than 0.2 mm; contains any spikes larger than 5mm. Following this filter, subjects who contained all four scans were retained. At the time of their first scan, the average subject age was 29.36 ± 3.36 years, with a range from 22 – 36. 70 of these subjects were monozygotic twins, 57 where non-monozygotic twins, and 2 were not twins.

A comprehensive description of the imaging parameters and image preprocessing can be found in [105] and in HCP’s online documentation (https://www.humanconnectome.org/study/hcp-young-adult/document/1200-subjects-data-release). T1w were collected on a 3T Siemens Connectome Skyra scanner with a 32-channel head coil. Subjects underwent two T1-weighted structural scans, which were averaged for each subject (TR = 2400 ms, TE = 2.14 ms, flip angle = 8°, 0.7 mm isotropic voxel resolution). fMRI were collected on a 7T Siemens Magnetom scanner with a 32-channel head coil. All 7T fMRI data was acquired with a gradient-echo planar imaging sequence (TR = 1000 ms, TE = 22.2 ms, flip angle = 45°, 1.6 mm isotropic voxel resolution, multi-band factor = 5, image acceleration factor = 2, partial Fourier sample = 7/8, echo spacing = 0.64 ms, bandwidth = 1924 Hz/Px). Four resting state data runs were collected, each lasting 15 minutes (frames = 900), with eyes open and instructions to fixate on a cross. Four movie watching data runs were collected, each lasting approximately 15 minutes (frames = 921, 918, 915, 901), with subjects passively viewing visual and audio presentations of movie scenes. Movies consisted of both freely available independent films covered by Creative Commons licensing and Hollywood movies prepared for analysis [114]. For both resting state and movie watching data, two runs were acquired with posterior-to-anterior phase encoding direction and two runs were acquired with anterior-to-posterior phase encoding direction.

Structural and functional images were minimally preprocessed according to the description provided in [105], as implemented and shared by the Connectome Coordination Facility. Briefly, T1w images were aligned to MNI space before undergoing FreeSurfer’s (version 5.3) cortical reconstruction workflow, as part of the HCP Pipeline’s PreFreeSurfer, FreeSurfer, and PostFreeSurfer steps. 7T fMRI images were downloaded after correction and reprocessing announced by the HCP consortium in April, 2018. fMRI images were corrected for gradient distortion, susceptibility distortion, and motion, and then aligned to the corresponding T1w with one spline interpolation step. This volume was further corrected for intensity bias and normalized to a mean of 10000. This volume was then projected to the 2mm *32k_fs_LR* mesh, excluding outliers, and aligned to a common space using a multi-modal surface registration [106]. The resultant cifti file for each HCP subject used in this study followed the file naming pattern: `*_Atlas_MSMAll_hp2000_clean.dtseries.nii`. These steps are performed as part of the HCP Pipeline’s fMRIVolume and fMRISurface steps. Resting state and moving watching fMRI images were nuisance regressed in the same manner. Each minimally preprocessed fMRI was linearly detrended, band-pass filtered (0.008-0.25 Hz), confound regressed and standardized using Nilearn’s `signal.clean` function, which removes confounds orthogonally to the temporal filters. The confound regression strategy included six motion estimates, mean signal from a white matter, cerebrospinal fluid, and whole brain mask, derivatives of these previous nine regressors, and squares of these 18 terms. Spike regressors were not applied. Following these preprocessing operations, the mean signal was taken at each time frame for each node, as defined by the Schaefer 400 parcellation [48] in *32k_fs_LR* space.

### Datasets: Nathan Kline Institute, Enhanced Rockland Sample 3T resting-state and diffusion weighted MRI

The Nathan Kline Institute Rockland Sample (NKI) dataset consisted of structural magnetic resonance imaging, resting state functional magnetic resonance imaging data, as well as diffusion magnetic resonance imaging data from 811 subjects (downloaded December 2016 from the INDI S3 Bucket) of a community sample of participants across the lifespan [65]. After excluding subjects based on data and metadata completeness and quality control, the final subset utilized included 542 subjects (56% female, age range = 7-84). The study was approved by the Nathan Kline Institute Institutional Review Board and Montclair State University Institutional Review Board and informed consent was obtained from all subjects. A comprehensive description of the imaging parameters can be found online at the NKI website.

Briefly, images were collected on a Siemens Magneton Trio with a 12-channel head coil. Subjects underwent one T1-weighted structural scan (TR = 1900 ms, TE = 2.52 ms, flip angle = 9°, 1 mm isotropic voxel resolution). Subjects underwent three differently parameterized resting state scans, but only one acquisition is used in the present study. The fMRI data was acquired with a gradient-echo planar imaging sequence (TR = 645 ms, TE = 30 ms, flip angle = 60°, 3 mm isotropic voxel resolution, multiband factor = 4). This resting state run lasted approximately 9:41 seconds, with eyes open and instructions to fixate on a cross. Subjects underwent one diffusion MRI scan (TR = 2400 ms, TE = 85 ms, flip angle = 90°, 2 mm isotropic voxel resolution, 128 diffusion weighted volumes, b-value = 1500 s/mm^2^, 9 b = 0 volumes).

The NKI was downloaded in December of 2016 from the INDI S3 Bucket. At the time of download, the dataset consisted of 957 T1w (811 subjects), 914 DWI (771 subjects), and 718 fMRI (“acquisition645”; 634 subjects) images. T1w and DWI images, and tractography results were first filtered based on visual inspection. T1w images were filtered based on artifact, such as ringing or ghosting (43 images) and for FreeSurfer reconstruction failure (105 images) as assesses with the ENIGMA QC tools, leaving 809 T1w images (699 subjects). DWI images were filtered based on corrupt data (13 images) and artifact on fitted fractional anisotropy maps (18 images), leaving 883 images (747 subjects). Tractography was run on 781 images (677 subjects) that had both quality controlled T1w and DWI images. Tractography results were filtered based on artifact, which include failure to resolve callosal, cingulum, and/or corticospinal streamlines or errors resulting in visually sparse streamline densities, resulting in 764 tractography runs (661 subjects). T1w, DWI, and fMRI images were then filtered using computed image quality metrics [115–117]. T1w images were excluded if the scan was marked as an outlier (1.5x the inter-quartile range in the adverse direction) in three or more of following quality metric distributions: coefficient of joint variation, contrast-to-noise ratio, signal-to-noise ratio, Dietrich’s SNR, FBER, and EFC. DWI images were excluded if the percent of signal outliers, determined by eddy_qc, was greater than 15%. Furthermore, DWI were excluded if the scan was marked as an outlier (1.5x the inter-quartile range in the adverse direction) in two or more of following quality metric distributions: temporal signal-to-noise ratio, mean voxel intensity outlier count, or max voxel intensity outlier count. fMRI images were excluded if greater than 15% of time frames exceeded 0.5mm framewise displacement. Furthermore, fMRI images were excluded the scan was marked as an outlier (1.5x the inter-quartile range in the adverse direction) in 3 or more of the following quality metric distributions: DVARS standard deviation, DVARS voxelwise standard deviation, temporal signal-to-noise ratio, framewise displacement mean, AFNI’s outlier ratio, and AFNI’s quality index. This image quality metric filtering excluded zero T1w images, 16 DWI images, and 21 fMRI images. Following this visual and image quality metric filtering, 809 T1w images (699 subjects), 728 DWI images (619 subjects), and 697 fMRI images (633 subjects). The intersection of subjects with at least one valid T1w, DWI, and fMRI images totaled 567 subjects. Finally, age metadata was available for 542 of these subjects.

T1-weighted images were submitted to FreeSurfer’s cortical reconstruction workflow (version 6.0). The FreeSurfer results were used to skull strip the T1w, which was subsequently aligned to MNI space with 6 degrees of freedom. fMRI preprocessing was performed using the fMRIPrep version 1.1.8 [118]. The following description of fMRI preprocessing is based on fMRIPrep’s documentation. This workflow utilizes ANTs (2.1.0), FSL (5.0.9), AFNI (16.2.07), FreeSurfer (6.0.1), nipype [119], and nilearn [120]. Each T1w was corrected using `N4BiasFieldCorrection` [107] and skull-stripped using `antsBrainExtraction.sh` (using the OASIS template). The ANTs derived brain mask was refined with a custom variation of the method to reconcile ANTs-derived and FreeSurfer-derived segmentations of the cortical graymatter of Mindboggle [121]. Brain tissue segmentation of cerebrospinal fluid (CSF), white-matter (WM) and gray-matter (GM) was performed on the brain-extracted T1w using `fast` [110]. Functional data was slice time corrected using 3dTshift from AFNI and motion corrected using FSL’s `mcflirt`. “Fieldmap-less” distortion correction was performed by co-registering the functional image to the same-subject T1w with intensity inverted [122] constrained with an average fieldmap template [123], implemented with `antsRegistration`. This was followed by co-registration to the corresponding T1w using boundary-based registration [124] with 9 degrees of freedom, using `bbregister`. Motion correcting transformations, field distortion correcting warp, and BOLD-to-T1w transformation warp were concatenated and applied in a single step using `antsApplyTransforms` using Lanczos interpolation. Frame-wise displacement [125] was calculated for each functional run using the implementation of Nipype. The first four frames of the BOLD data in the T1w space were discarded. Diffusion images were preprocessed following the “DESIGNER” pipeline using MRTrix (3.0) [126, 127], which includes denoising, Gibbs ringing and Rician bias correction, distortion and eddy current correction [128] and B1 field correction. DWI were then aligned to their corresponding T1w and the MNI space in one interpolation step with B-vectors rotated accordingly. Local models of white matter orientation were estimated in a recursive manner [129] using constrained spherical deconvolution [109] with a spherical harmonics order of 8. Tractography was performed using Dipy’s `Local Tracking` module [108]. Probabilistic streamline tractography was seeded five times in each white matter voxel. Streamlines were propagated with a 0.5 mm step size and a maximum turning angle set to 20°. Streamlines were retained if longer than 10 mm and with valid endpoints, following Dipy’s implementation of anatomically constrained tractography [112]. Streamline count between regions-of-interest were normalized by dividing the count between regions by the geometric average volume of the regions.

#### Estimating group-representative structural connectivity network

The output of the tractography algorithm generated subject-level estimates of streamlines for both the NKI and HCP datasets. In general, subjects’ connectomes are variable. A fraction of this variability reflects true individual differences, while another fraction reflects unwanted noise, e.g. random variation. One strategy for reducing noise is to aggregate data from many individuals to construct a group-representative consensus connectome. Here, we follow [44] and generate distant-dependent connectomes for both the NKI and HCP datasets. Briefly, this procedure bins edges by their length and, within each distance bin, identifies the edges that are most consistently present across the full set of subjects. Compared to standard approaches, which retains the most consistent edges irrespective of their length, consensus networks generated using this procedure are more representative of single-subject connectomes–i.e. has more properties in common. Note that this distance-preserving consensus procedure is applied separately to within- and between-hemisphere edges. Note also that for the NKI dataset, the consensus connectome was constructed using data from subjects aged 18-35 years. Finally, we made the decision to use the same (but dataset-specific) group-representative for all HCP and NKI subjects. The rationale behind this decision was that it allowed us to discount the possibility that differences in model performance–e.g. fitness or edge weights–was driven by differences in the structural connectivity.

### Dataset: Mouse anatomical and functional connectivity

#### Mouse resting state fMRI data

All in vivo experiments were conducted in accordance with the Italian law (DL 2006/2014, EU 63/2010, Ministero della Sanitá, Roma) and the recommendations in the Guide for the Care and Use of Laboratory Animals of the National Institutes of Health. Animal research protocols were reviewed and consented by the animal care committee of the Italian Institute of Technology and Italian Ministry of Health. The rsfMRI dataset used in this work consists of n = 19 scans in adult male C57BL/6J mice that are publicly available [130, 131]. Animal preparation, image data acquisition, and image data preprocessing for rsfMRI data have been described in greater detail elsewhere [131]. Briefly, rsfMRI data were acquired on a 7.0-T scanner (Bruker BioSpin, Ettlingen) equipped with BGA-9 gradient set, using a 72-mm birdcage transmit coil, and a four-channel solenoid coil for signal reception. Single-shot BOLD echo planar imaging time series were acquired using an echo planar imaging sequence with the following parameters: repetition time/echo time, 1000/15 ms; flip angle, 30°; matrix, 100 ×100; field of view, 2 × 2 cm2; 18 coronal slices; slice thickness, 0.50 mm; 500 (n = 21) or 1500 (n = 19) volumes; and a total rsfMRI acquisition time of 30 min.

Image preprocessing has been previously described in greater detail elsewhere [131]. Briefly, timeseries were despiked, motion corrected, skull stripped and spatially registered to an in-house EPI-based mouse brain template. Denoising and motion correction strategies involved the regression of mean ventricular signal plus 6 motion parameters. The resulting time series were band-pass filtered (0.01-0.1 Hz band) and then spatially smoothed with a Gaussian kernel of 0.5 mm full width at half maximum. After preprocessing, mean regional time-series were extracted for 182 regions of interest (ROIs) derived from a predefined anatomical parcellation of the Allen Brain Institute (ABI, [13, 132]).

#### Mouse Anatomical Connectivity Data

The mouse anatomical connectivity data used in this work were derived from a voxel-scale model of the mouse connectome made available by the Allen Brain Institute [133, 134] (https://data.mendeley.com/datasets/dxtzpvv83k/2).

Briefly, the mouse structural connectome was obtained from imaging enhanced green fluorescent protein (eGFP)–labeled axonal projections derived 428 viral microinjection experiments, and registered to a common coordinate space [13]. Under the assumption that structural connectivity varies smoothly across major brain divisions, the connectivity at each voxel was modeled as a radial basis kernel-weighted average of the projection patterns of nearby injections [134]. Following the procedure outlined in [133], we re-parcellated the voxel scale model in the same 182 nodes used for the resting state fMRI data, and we adopted normalized connection density (NCD) for defining connectome edges, as this normalization has been shown to be less affected by regional volume than other absolute and/or relative measure of interregional connectivity [135].

### Fitting edge weights

Here, we use a regression-based framework for assigning weights to existing structural connections. Our approach is simple; we assume that at time *t* the state of region *i* (level of fMRI BOLD activity) is a function of its neighbors’ states at time *t* – 1 plus an offset (bias). That is:

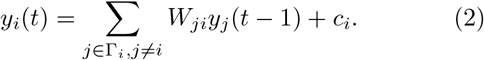

Here, *y*i**(*t*) refers to the level of activity in region *i* at time *t*, Γ_i_ is the set of *i*’s connected neighbors (their indices). We use linear regression and ordinary least squares to estimate the parameters *W_ji_* and *c_i_* separately for each node *i*. Thus, the resulting matrix 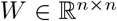, is sparse and preserves *exactly* the binary structure of the whitematter connectivity. However, the weights can take on both positive and negative valence. The resulting network is also asymmetric–i.e. in general, the *W_ij_* ≠ *W_ji_*.

### Null models

We fit the linear model using data pooled from all participants and scans. The model fitness was quantified as the mean squared error (MSE) of the observed activity time series and the time series predicted by the model. We compared the empirical MSE against five null models.

- *Minimally wired null model*: Generates a synthetic structural network comprised of the *m* least costly connections, where *m* is the same number of connections as the observed network. Because this network contains only short-range (low cost) connections, this null model assesses how long-distance connections contribute to model fitness.
- *Re-ordered null model:* Randomly permutes node order, effectively endowing nodes with a different number and set of neighbors than they have in the original network. This model assesses the contributions of specific neighbors to model fitness.
- *“Spin” null model:* Randomly permutes node order while approximately preserving inter-regional Euclidean distances. This model can be viewed as a constrained versions of the *re-ordered null model,* in that it only allows particular subset of permutations.
- *Topological null model:* In this model, each node makes the same number of connections as in the original network. However, those connections, which define nodes’ neighborhoods, are formed at random. This model assesses whether networks with identical degree distribution yield similar fitness values as the original network.
- *Temporal null model:* For each scan, parcel time series are circularly shifted by some random integer. This procedure preserves temporally invariant properties of each time series, like their mean and standard deviation, and approximately preserves other properties, e.g. power spectrum. However, it destroys interregional correlations. In effect, this model tests whether time series with similar statistical properties but no correlation structure could yield comparable fitness values as the original time series.

### Modularity maximization

Here, we used modularity maximization to detect clusters (modules) in brain network data [136, 137]. Generically, modularity maximization works by assigning nodes to non-overlapping clusters so that the within-cluster weight of connections maximally exceeds that of a null model. This intuition is formalized by the modularity quality function:

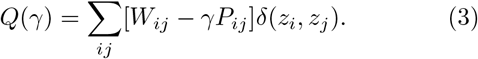

In this equation, *W_ij_* and *P_ij_* are the observed and expected weight of the connection between nodes *i* and *j*, *z_i_* ∈ {1,…, *K*} is a categorical variable that indicates the community to which node *i* was assigned, *γ* is the structural resolution parameter, and *δ*(*z_i_*, *z_j_*) is the Kronecker delta function, which evaluates to 1 when *z_i_* = *z_j_* and 0 otherwise. In short, modularity maximization seeks to optimize the quantity *Q*(*γ*) by selecting the values of *z_i_*.

The modularity maximization framework is general and can test different null hypotheses (null connectivity models) by varying the entries of *P*, the matrix of expected connections and their weights. Here, we test two different null models. The first was proposed in [51] and is designed, specifically, to work with signed connectivity matrices. Under this model, the modularity equation is:

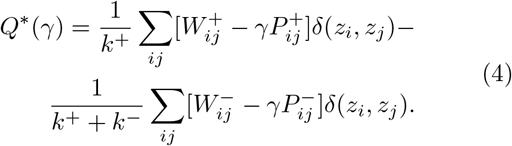

Here, the modularity equation includes separate terms for the positive and negative connections. The positive term is weighted more than the negative term (note the scale factors before the summation). This allows modules to be detected in networks with signed connections. However, if this same version of modularity maximization is applied to a network with positive links only, the second term in the equation evaluates to zero and returns the standard modularity equation. Note that in this equation, 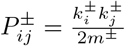.

Here, we use this quality function in two ways. In the main text, we optimize *Q*\* 1000 times for both the asymmetric, weighted, and signed network as well as the fiber density network. These results are shown in Fig. 2. In the supplement, we combine this quality function with a hierarchical consensus algorithm [54], in which we first vary the values of *γ* over all possible ranges to obtain a representative sample of communities (1,000,000 repetitions in total), and second, use these samples to construct a hierarchical dendrogram that organizes the noisy individual samples into hierarchically related consensus communities. The results of this analysis are shown in Fig. S9.

We also used a second version of the modularity equation that was originally proposed for analysis of physical systems [59]. Briefly, the equation reads:

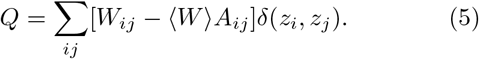

Here, the matrix *A* is the binary matrix of connections that exist in the empirical and weighted connectivity matrix. 〈*W*〉 is the mean weight of existing connections. In other words, this modularity equation preserves the topology of the network, but assumes that edge weights are assigned randomly and uniformly. The results of this analysis are presented in Fig. S11. We note that, in principle, a resolution parameter could be incorporated into this formulation of the modularity quality function as well by replacing 〈W〉 with a tunable *γ* parameter.

### Network statistics

In addition to modularity, we calculated several other network metrics. These include efficiency, characteristic path length, signed strength, and signed clustering coefficient. In this section we define those measures in detail.

- **Shortest paths.** Both efficiency and characteristic path length are defined based on a network’s shortest path structure. Consider source and target nodes, *s* and *t*. The shortest path from the *s* to *t* can be estimated easily using the Floyd-Warshall algorithm. In weighted networks where edge weights as interpreted as measures of affinities it requires that the user first map those weights to measures of cost. For networks with positive connections only, a straightforward way to do this is to transform 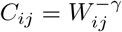, where the most common value for the parameter is *γ* = 1. For signed networks, like the ones used here, we use the same transformation, but only after we add an offset to each edge so that all weights are greater than zero. Our strategy for doing so involved first subtract the smallest (most negative) edge weight from the network. This ensures that all edges have a weight greater than zero, except for the single edge corresponding to the most negative weight, which has a cost of 0. We then add to every edge an even smaller offset–in this case the weakest edge weight in the fiber density matrix. This guarantees that all pairs of nodes connected by a fiber tract have nonzero weights. Once a network’s affinity-based weights have been transformed to costs, algorithms like the Floyd-Warshall algorithm find the shortest–i.e. least costly–path between all pairs of nodes. This algorithm returns two outputs: 1) the total cost incurred by following said path and 2) the number of steps (hops) along said path. Here, we use the hop data but not that, in principle, one could repeat all subsequent analyses using the cost data, instead. Let *H_st_* be the number of hops on the shortest path from the source s to the target *t*.
- **Characteristic path length.** The characteristic path length of this network is calculated as:

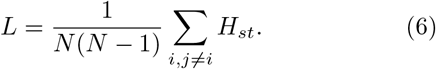
- **Efficiency.** The efficiency of this network is calculated as:

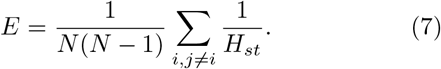
- **Clustering coefficient.** The local clustering coefficient is calculated for each node *i*. Intuitively, it measures the extent to which node *i*’s neighbors are also connected to one another. It can be calculated easily for each node as the density of the subgraph composed of those neighbors. Here, we calculate clustering coefficients for each node in the network based on their positive connections and negative connections, separately. The values reported in the main text ignore the actual weight but preserve sign.
- **Strength.** Node strength – or weighted degree – the total weight of connections incident upon node *i*. For an undirected network, it is calculated as: *s_i_* ∑_*j*_ *W_ij_*. For a directed network, we calculate strength as the average of a nodes’ incoming and outgoing connections, i.e. 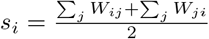. Here, we also differentiate between a node’s positive and negative strength. Let *W*^+^ and *W*^-^ be the networks of positive and negative connections only. For the network of negative connections, we conveniently flip the sign of each connection. Then we calculate each nodes’ signed strength as 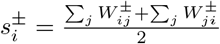
- **Partition laterality.** We calculated partition laterality following Lohse *et al.* [53]. For a given community *c*, we calculate its uncorrected laterality as 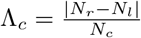. Here, *N_c_* is the number of nodes in *c* and *N_r_* and *N_l_* are the number of those nodes in the right and left hemispheres, respectively. When the community has a balanced number of nodes from both hemispheres its laterality is close to zero; if it is left- or right-dominant, then the value is close to 1. For a partition comprised of communities *c*_1_,…, *c_K_*, we calculate the partition laterality as 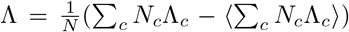. Here, the term ∑_*c*_ *N_c_*Λ_*c*_) indicates the expected laterality under a null model in which nodes get randomly assigned to one hemisphere or another. Note that here we cannot use spin tests for the permutation; the spin tests preserves hemisphere labels and a “spun” partition would have laterality exactly equal to that of the original, unpermuted partition.

### Neural mass models

Many studies have tried to link brain structure and function [49]. One popular strategy for doing so is to use an estimate of anatomical connectivity to generate synthetic covariance matrices (either directly or by first generating synthetic neural time series and calculating their covariance empirically). The synthetic covariance matrices can then be compared to the empirical FC, usually as a correlation of their edge weights. The resulting coefficient serves as a measure of structure-function coupling. Here, we analyzed two models for generating synthetic covariance matrices or time series based on population-level “neural mass” models (NMMs).

- **Galan model.** We follow work by Honey *et al.* [77] for estimating the inter-areal covariance matrix, **C**, based on a linearization of Wilson-Cowan dynamics for neuronal populations [138]. The element *C_ij_* ∈ **C** denotes the covariance of activity in area *i* with that of area *j*. In more detail, we let **u**(*t*) = {*u*_1_(*t*),…, *u_N_*(*t*)} be the vector of brain areas’ states (activity levels) at time *t*. Under these dynamics, brain areas’ states evolve as:

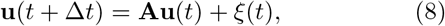

where *ξ*(*t*) is uncorrelated Gaussian noise and Δ*t* is a single time step. Here, the generalized coupling matrix, **A**, is based on the structural connectivity matrix, **W**, and was defined as:

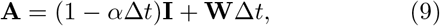

where *α* is a leak variable within each brain area and **I** is the identity matrix. As in Honey *et al.* [77], we fixed *α* = 2. Conveniently, Galán [138] showed that brain areas’ pairwise covariances (summarized by the matrix 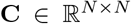) can be estimated directly from the spectral properties of **A** and the covariance of the noise terms *ξ*(*t*). As with covariance matrices estimated from recorded time series of brain activity, we interpret **C** as an estimate of functional connectivity. See Galán [138] for more details.
- **Reduced Wong-Wang mean field model.** We also studied a second biophysical model for fMRI BOLD data. Unlike the Galán model, which calculates the covariance structure analytically given a structural connectivity matrix, this model generates simulated time series, first by using a reduced spiking neural network to generate population-level time courses, and second by convolving these data with a hemodynamic model. The spiking network model evolves according to the following differential equations:

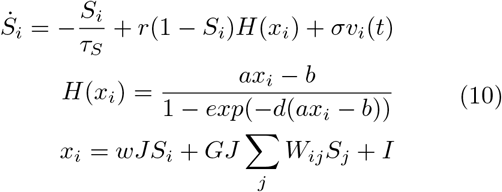 In this equation, *x_i_*, *H*(*x_i_*), and *S_i_* are the total input current, population firing rate, and synaptic gating for region *i*. The input current, *x_i_* depends on recurrent connection strength, *w*, excitatory input, *I*, and inter-regional information “flow”, which is calculated as the sum of region *i*’s connected neighbors’ synaptic gating, weighted by the global coupling constant, *G*, and synaptic coupling constant, *J*. Following Wang *et al.* [139], we set the parameters of the input-output function, *H*(*x_i_*) to *a* = 270 n/C, *b* = 108 Hz, and *d* = 0.154 s. Kinetic parameters for synaptic activity were fixed at *r* = 0.641 and *τ_s_* = 0.1 s. The variable *v_i_*(*t*) is uncorrelated Gaussian-distributed noise whose variance is scaled by *σ*. This model generates neural activity at submillisecond timescales. Again, following Wang *et al.* [139], population level activity is input to the Balloon-Windkessel hemodynamic model [28], which yields simulated fMRI BOLD time courses for every brain region.

---

**FIG. S1.**
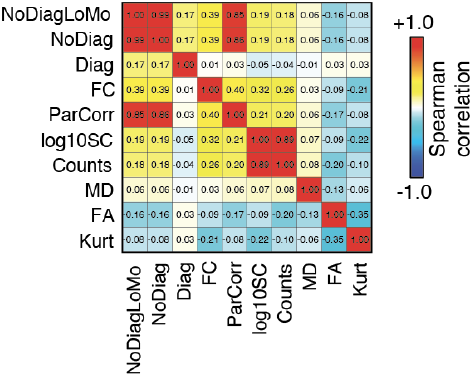
Similarity of edge weights. In the main text we use linear models to fit edge weights to structural connections. Here, we different edge weighting schemes. We consider: the logarithm of fiber densities (log10SC), estimated edge weights using all data (NoDiag), estimated edge weights using only low-motion data (NoDiagLoMo), estimated edge weights with the inclusion of an autoregressive diagonal term (Diag), functional connectivity estimated using a full correlation (FC), and partial correlations (ParCorr). In all cases, similarity was calculated as the Spearman correlation using only edges for which a structural connection was observed.

**FIG. S2.**
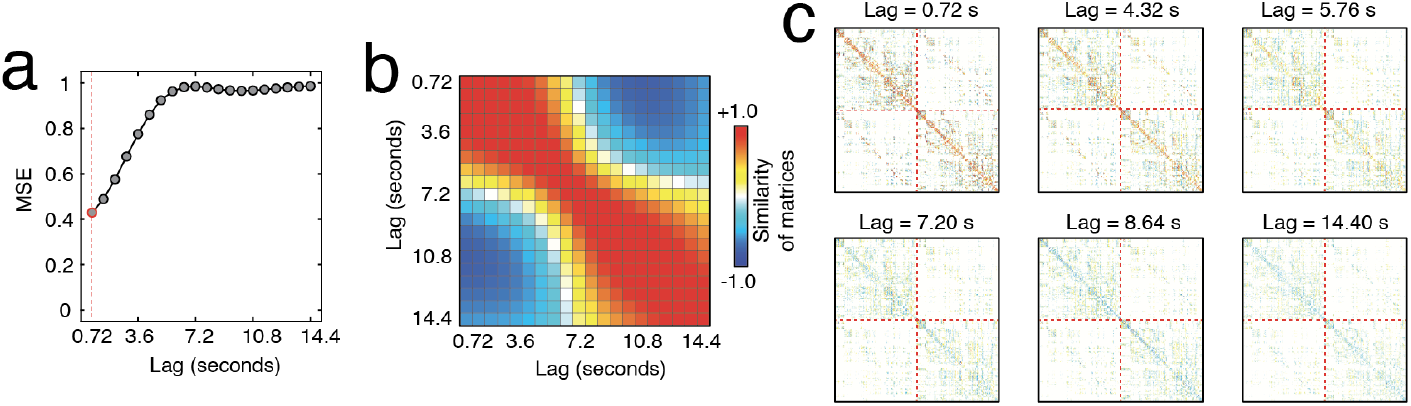
Effect of lag on estimated edge weights. In the main text we use linear models to fit edge weights by predicting activity at time *t* + 1 using activity at time *t* (one frame of lag = 0.720 seconds). Here, we investigate the effect of longer lags on the estimated weights, increasing the lag to ≈15 seconds. (*a*) Model error at different lags. (*b*) Spatial similarity of weights estimated at each lab. (*c*) Example weight matrices at different lags.

**FIG. S3.**
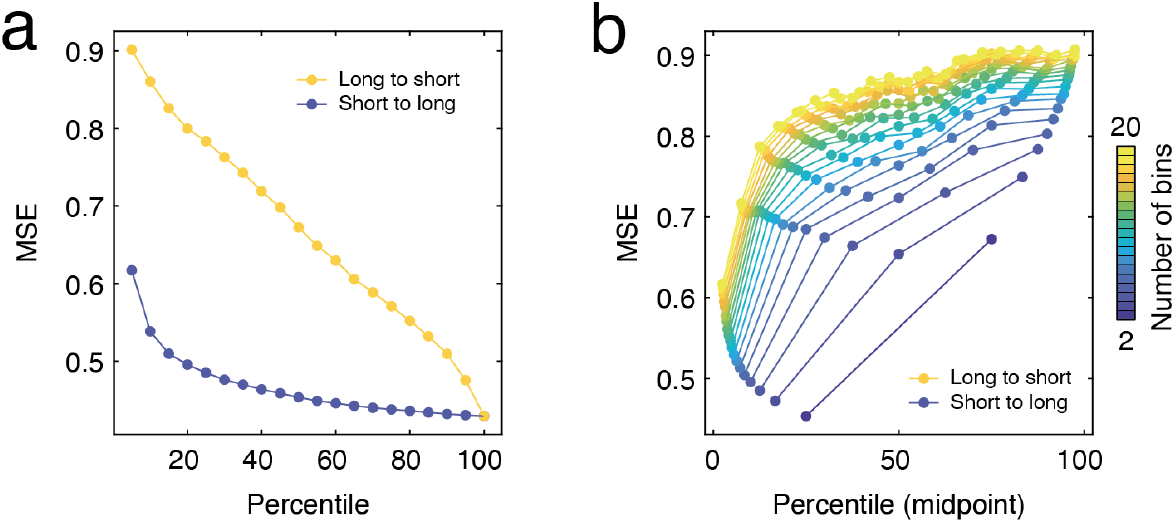
Effects of connection length on model performance. We divided connections into 20 percentile bins based on their length. Next, we created structural network backbone comprising the 5% shortest (or longest) connections, gradually adding back the progressively longer (or shorter) connection 19 bins, until all connections were added. This allowed us to assess the relative contributions of short *versus* long connections. In panel *a*, we plot the model error as a function across all percentiles. We also grouped connections by percentiles (from 2 to 20) and fit the model using each percentile independently rather cumulatively. The results in panel *b* suggest that short connections are more informative and lead to reduced model error.

**FIG. S4.**
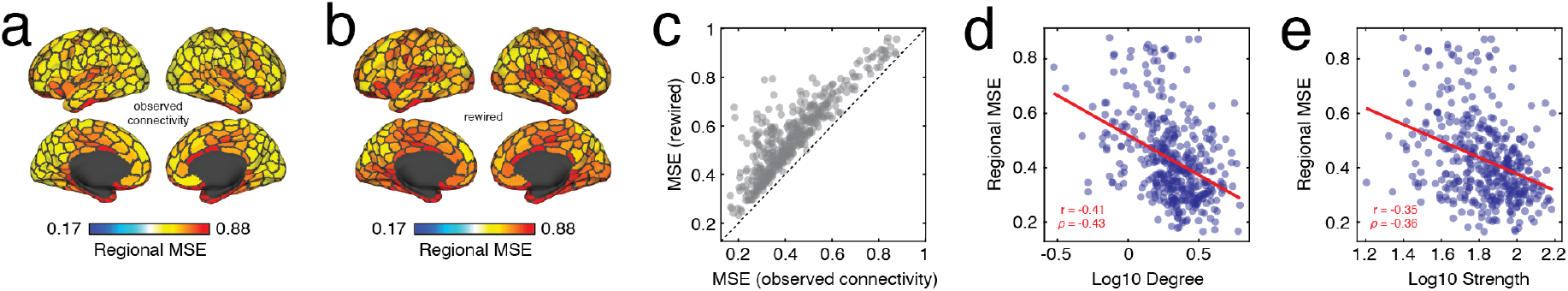
Model errors at the regional level. Regional mean squared error (MSE) for model fit using observed and intact network (*a*) *versus* the mean MSE across 100 models fit using a degree-preserving rewiring model. (*c*) Scatterplot comparing regional error. Points above the diagonal are regions fit better using the observed network compared to the rewired. Panels *d* and *e* compare regional MSE with the logarithm of nodes’ degree and weighted degree. That is, nodes with fewer connections exhibited worse fits (greater MSE).

**FIG. S5.**
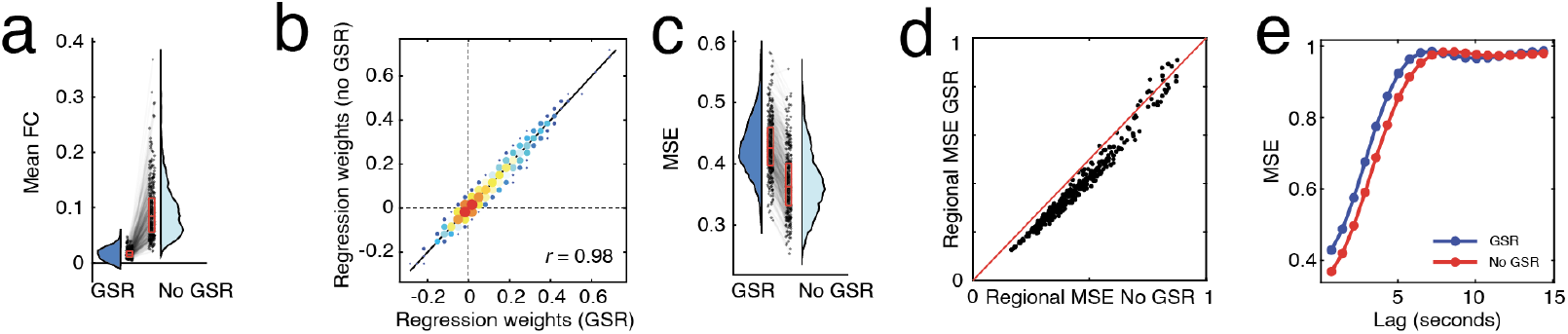
Effect of global signal regression on results. The human data analyzed in the main text was processed using a procedure that included global signal regression (GSR). Here, we compare select results using data processed without GSR. (*a*) Mean FC for GSR and non-GSR data; each point is a scan (4 scans × 95 participants yields 380 points). (*b*) Two-dimensional histogram of group-level edge weights estimated using GSR and non-GSR data. (*c*) Subject-level model error. (*d*) Comparison of regional model errors. The red line is identity (equal performance with both models). Points below the line have time series better predicted with non-GSR data; points above the line have activity time series better predicted by GSR data. (*e*) Model error as a function of lag.

**FIG. S6.**
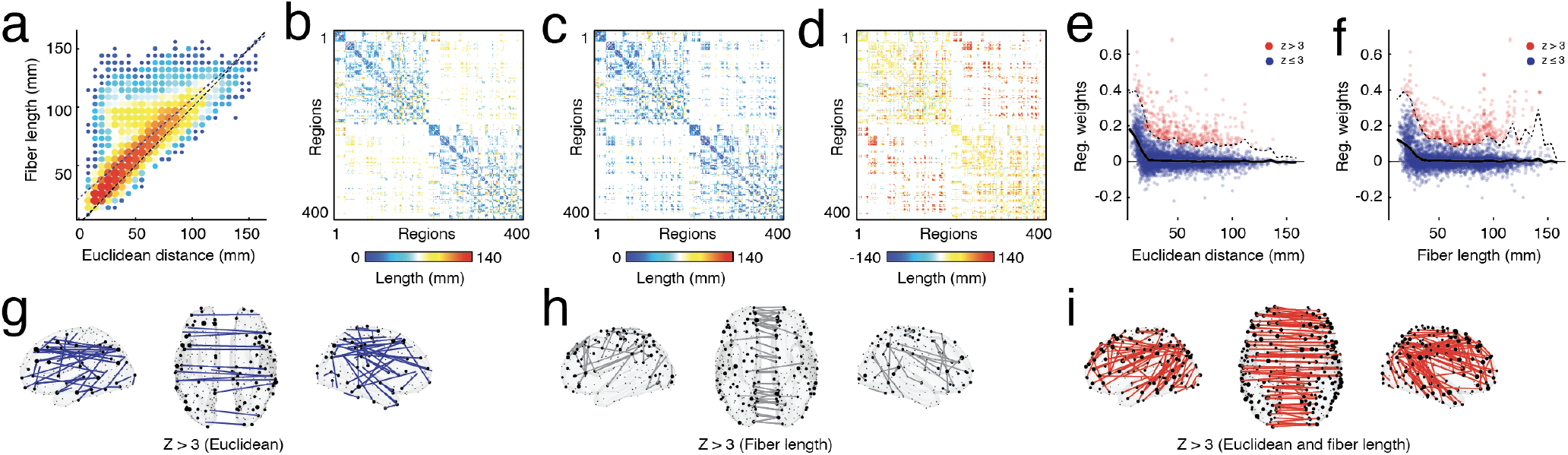
Dependence of distance effects on Euclidean distance *versus* fiber length. (*a*) Euclidean distance *versus* fiber length. Panels *b* and *c* depict edges weighted by fiber fiber length and Euclidean distance. (*d*) Difference in fiber length and Euclidean distance. Panels *e* and *f* show connection weight versus distance. Panels *g-i* show connections that are stronger than expected but unique to Euclidean distance, fiber length, or are shared between both.

**FIG. S7.**
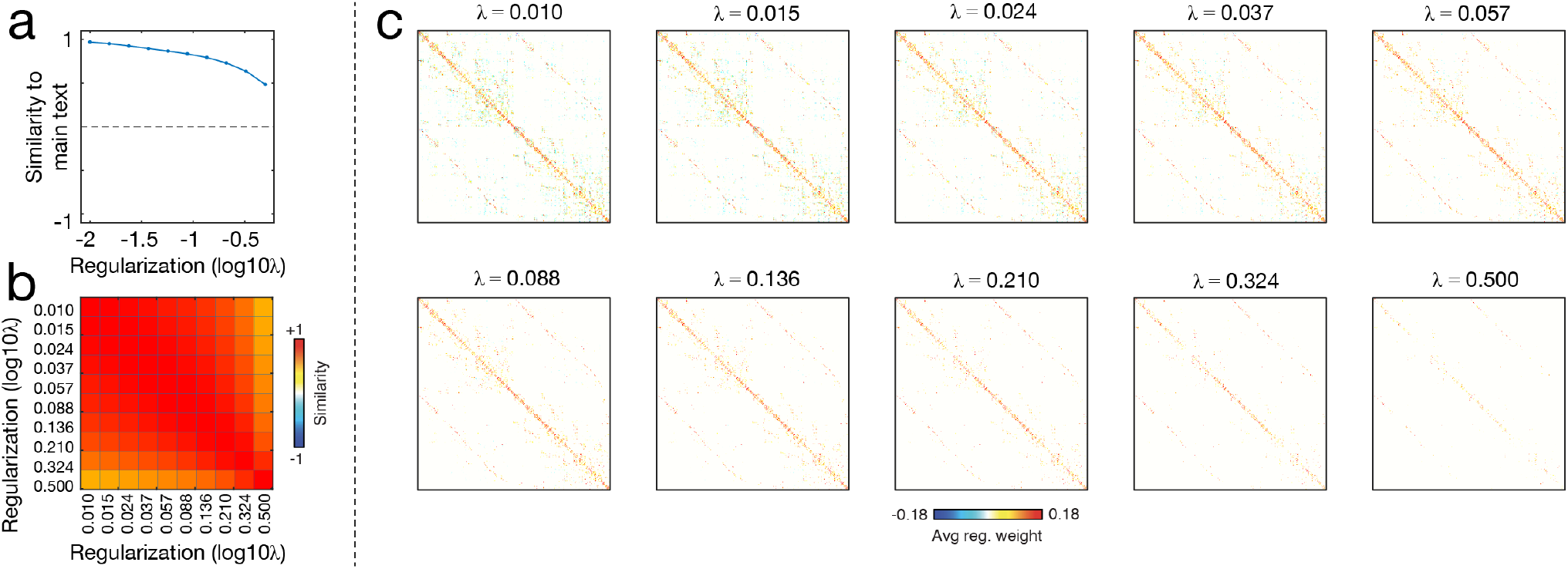
Comparing connectivity weights estimated with an without regularization. In the main text we used linear regression optimized with ordinary least squares to fit weights to edges. Here, we compare those results to weights fit using lasso regularization (implemented in MATLAB with the function `lasso.m`). For each node, we fits its edge weights independently. We logarithmically sampled 10 values of the regularization parameter, *λ*, over the interval [10^-3^,0.5]. For each value of *λ*, we generate a new estimate of connection weights, with sparsity increasing monotonically. (*a*) Correlation of edge weight values with the edge weights reported in the main text (no regularization). (*b*) Similarity of regularized connection weights to one another. (*c*) Connectivity matrices.

**FIG. S8.**
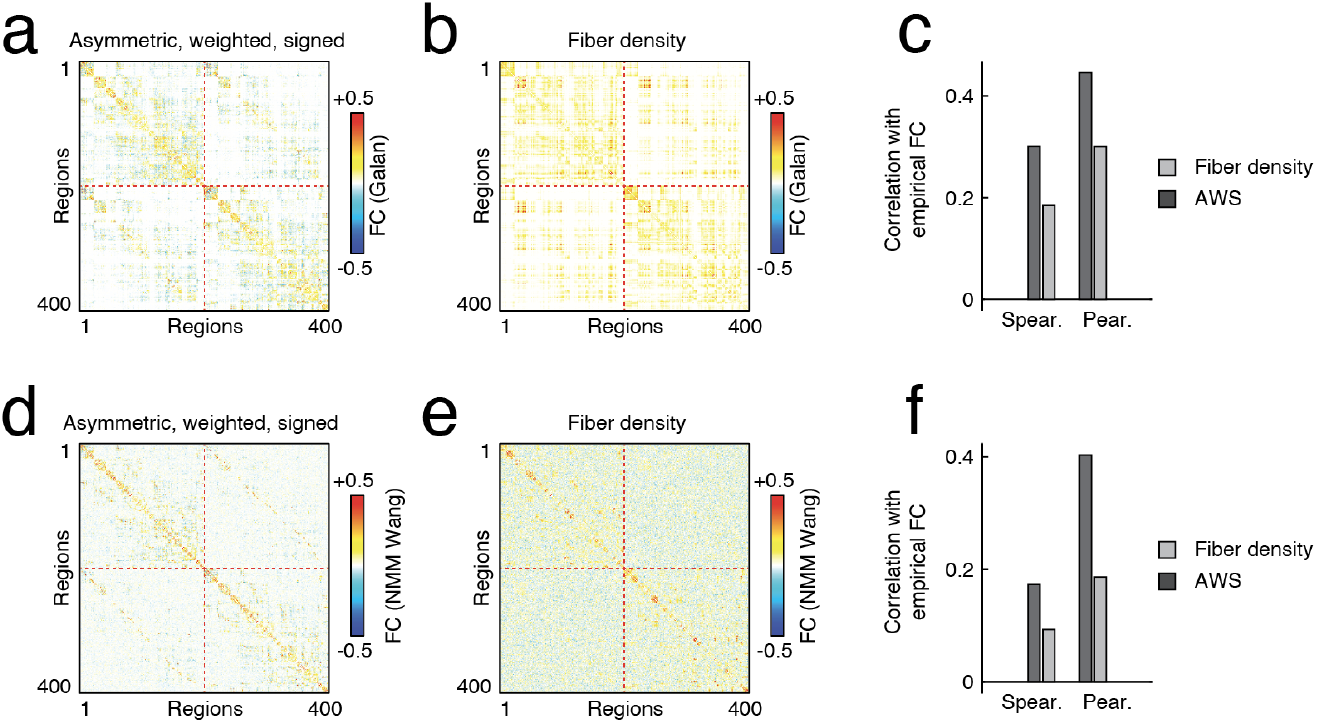
Simulated FC using linearized neural mass model. We used a linearization of a neural mass model [138] to generate simulated covariance matrices (which are then scaled to correlation matrices; FC). We performed this procedure using both the asymmetric, weighted, and signed version of SC as well as the traditional fiber density matrix. Panels *a* and *b* show the resulting correlation matrices. We then compared these matrices to an empirical estimate of FC. We found that the correlation matrix generated using asymmetric, weighted, and signed matrix was more similar to FC than the matrix generated using the fiber density weights. Panel *c* shows the results of this analysis using both product-moment correlation and rank correlations to assess similarity. We also analyzed a simplified neural mass model (reduced Wong-Wang model with implementation made available from [139]). This procedure generates synthetic fMRI BOLD time series that can be treated identically to an empirical time series. The correlation structure of the synthetic data can then be compared to the empirical correlation structure, i.e. FC. Panels *d-f* depict correlation matrices for asymmetric, weighted, and signed structural connectivity, the fiber density matrix, and their correlations with respect to the empirical FC matrix.

**FIG. S9.**
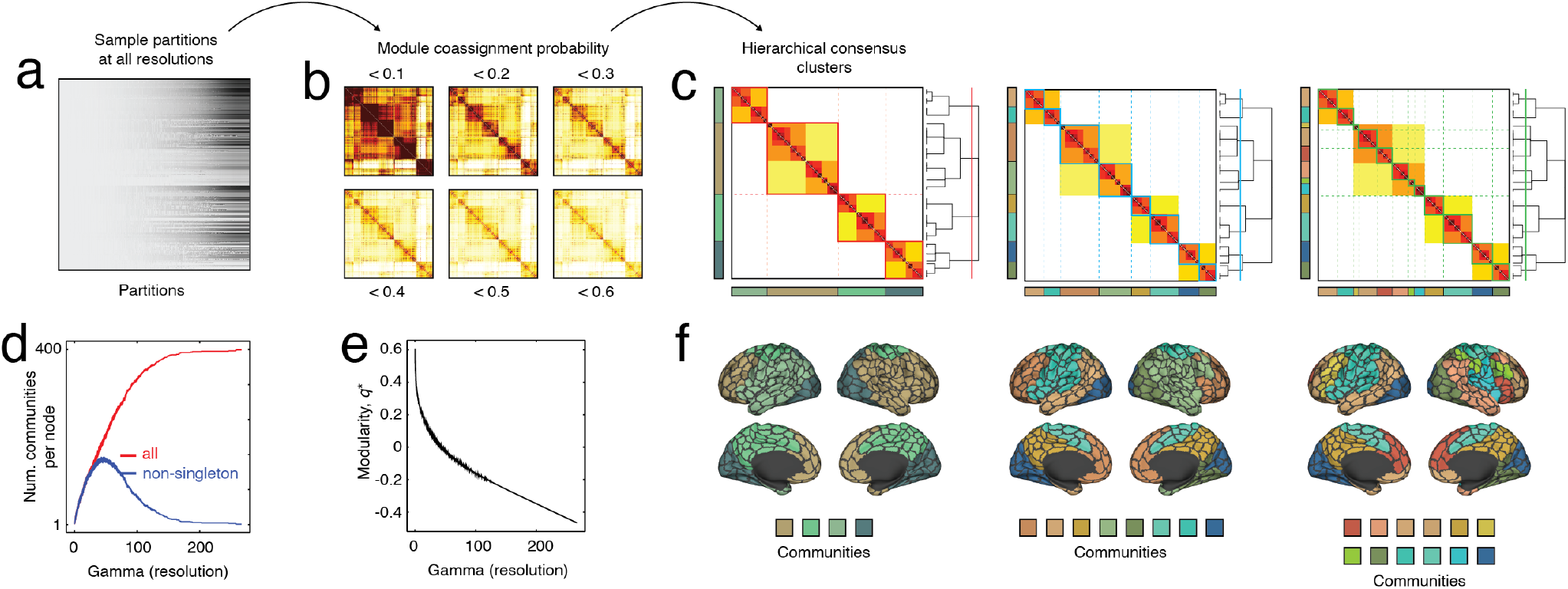
Comparing modular structure of structural networks. Modules are cohesive subnetworks – nodes that make more connections to other members of the same module than to other modules. Here, we compare the modular structure of network with weights defined by fiber density and the same network with asymmetric, weighted, and signed edges. Here, we examine modules estimated with a fixed resolution parameter (*γ* = 1) but explore the multiscale modular structure in the supplement. Co-assignment probability matrices for the inferred edge weight (*a*) and the fiber density matrices (*b*). (*c*) Element-wise difference in module co-assignment. (*d*) System co-assignment matrix for reference. Comparison of modularity (*e*) and laterality (*f*) of detected modules. (*g*) Alignment of modules with respect to coarse- and fine-system partitions. (*h* and *i*) Consensus communities for both versions of weights.

**FIG. S10.**
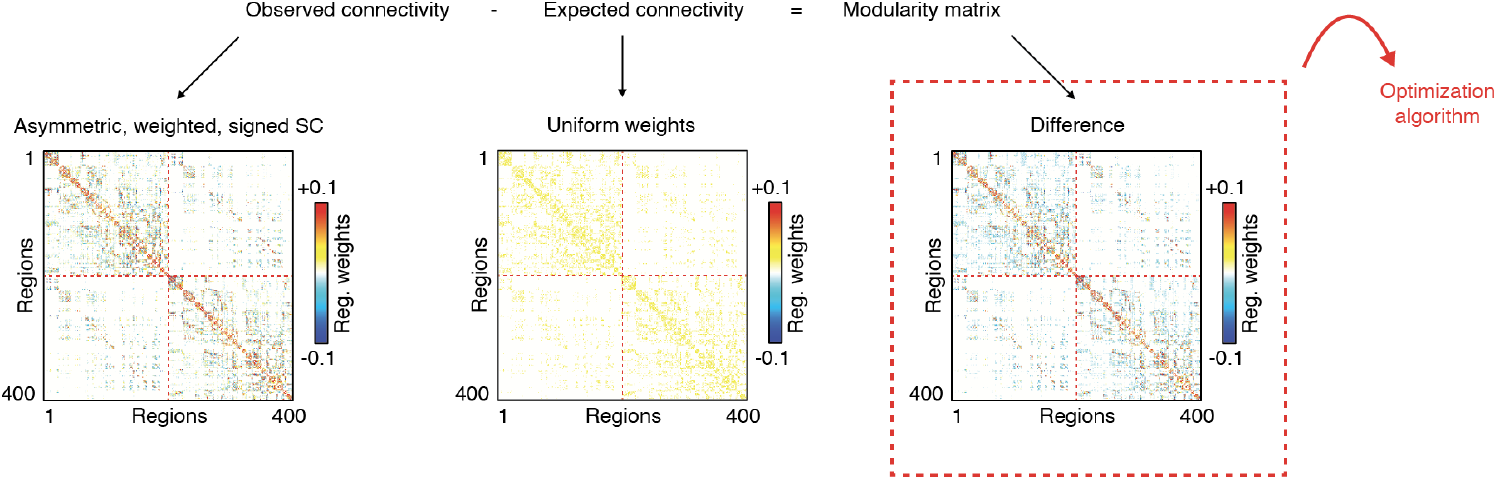
Modularity maximization with alternative, “geographic” null model. In the main text, we described the results of detecting modules using modularity maximization with an “standard” internal null model that preserved nodes’ (signed) degrees. Here, we report results using an alternative null model. Briefly, this null model preserves the same binary “backbone” of the original network, but assigns an expected weight to each connection equal to the mean weight of all existing connections. Modules therefore reflect groups of brain regions whose observed connections’ weights are greater than the mean weight.

**FIG. S11.**
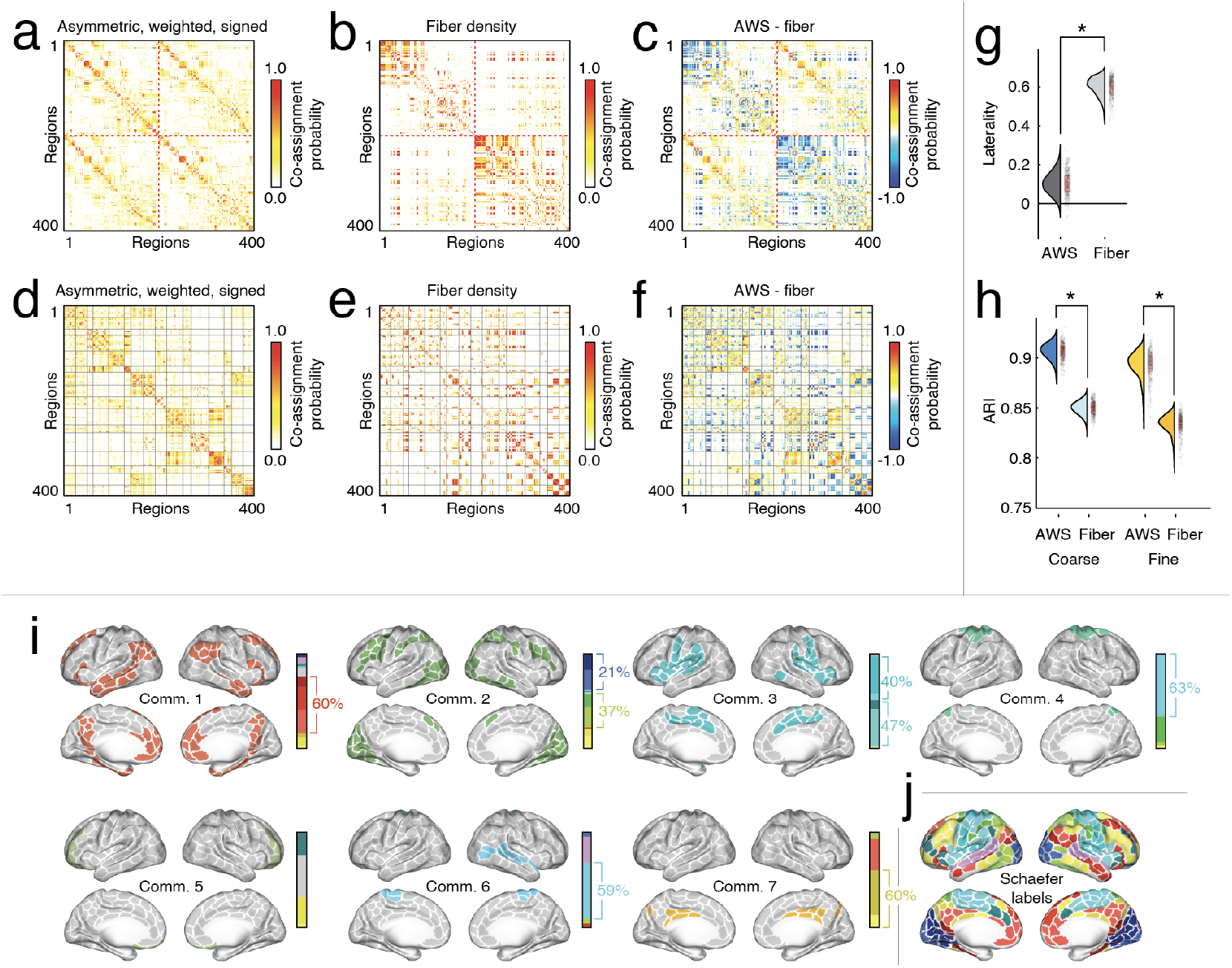
Modularity results using alternative null model. Module coassignment matrices for the signed, weighted, and directed SC (*a*) and fiber density (*b*). (*c*) Difference in coassignment probability. Panels *d-f* show the same matrices, but ordered by brain system. (*g*) Laterality of detected communities. Bilateral communities have values close to zero. (*h*) Adjusted Rand index of detected partitions with respect to coarse and fine-scale system labels. (*i*) Consensus communities detected in signed, weighted, and directed SC. The panel next to each surface plot depicts each community’s composition in terms of canonical brain systems. (*j*) Canonical brain systems.

**FIG. S12.**
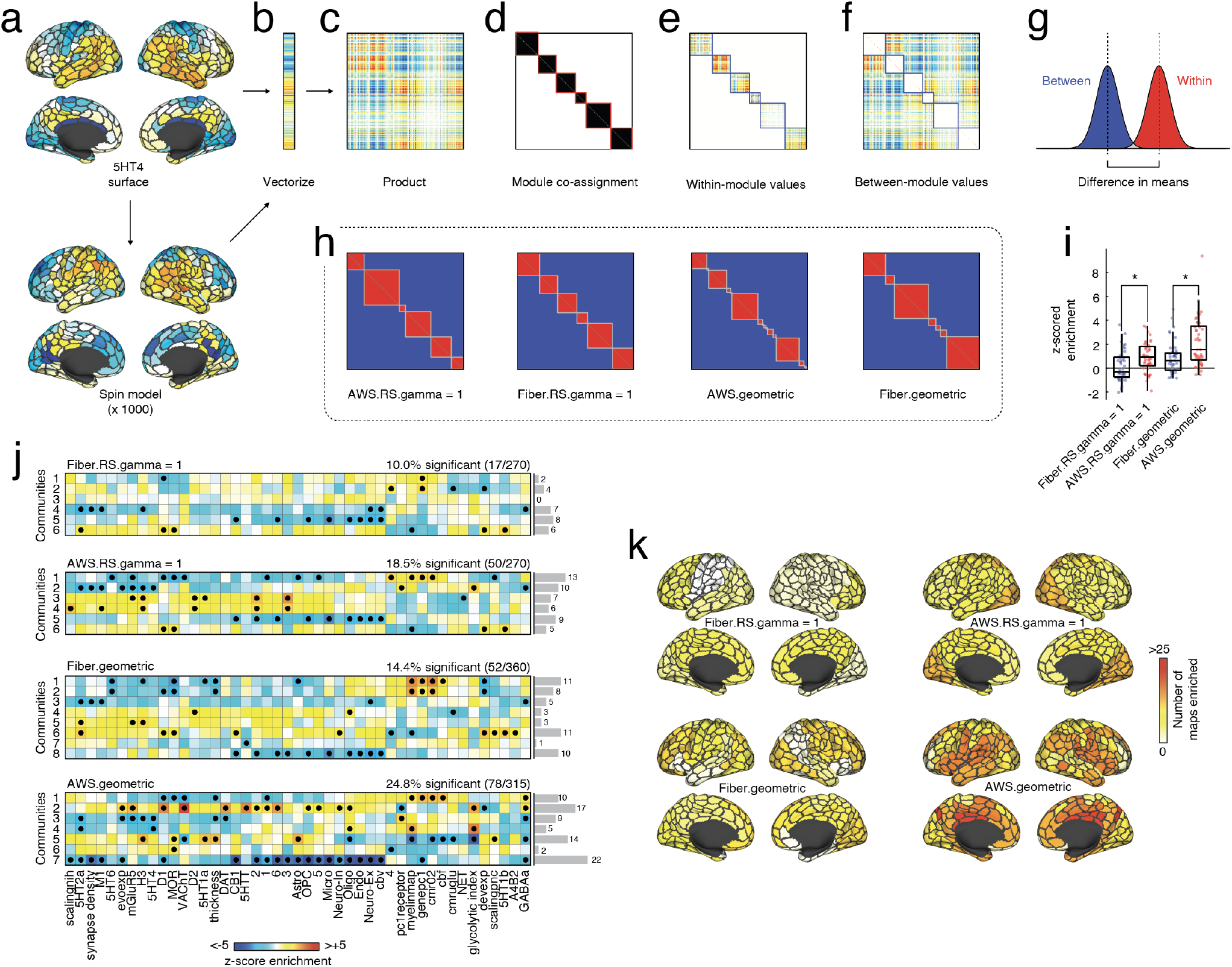
Enriching modules using brain maps. In the main text we performed community detection on the asymmetric, weighted, and signed network (`AWS`) and fiber density network (`Fiber`) using two versions of modularity maximization: one that uses a signed internal null model (RS; [51])and another that we termed a “geometric” null model, in which topology is preserved but weights are not (`geometric`; [59]). Here, we further explore, validate, and compare the modules by aligning them with respect to whole-brain maps made available as part of [60]. In total, we considered 45 maps that included receptor densities, gene expression, myelination status, evolutionary and developmental expansion, and synaptic density (among others). Here, we assess whether modules are “enriched” for these maps. That is, whether the module boundaries delineate spatial boundaries in each map. The procedure for doing so is illustrated using the serotonin receptor 5HTA as an example (see panel *a* for spatial distribution on cortical surface0. This map vectorized (*b*), and transformed into a matrix by calculating its outer product (*c*). We then compare the matrix to module co-assignment matrices (and example is shown in *d*). Specifically, we calculate the mean values of all within- and between-module elements (*e* and *f*) and subsequently calculate the difference in these means (*g*). In parallel, we repeat this procedure for 1000 spatially constrained permutations of the map (spin test). The spin tests generate a null distribution against which we compare the empirical (observed) difference in means *via* the z-score. Larger *z* values indicate greater modular “enrichment”. We perform this entire procedure for four sets of modules – signed and geometric model for both the asymmetric, weighted, and signed matrix as well as the fiber density matrix. Panel *i* shows the distribution of *z*-scores. In general, we find that the AWS model always outperforms the fiber density matrix–i.e. leads to greater z scores (paired sample *t*-test; maximum *p* = 0.0017). We also tested enrichment at the level of individual models (*j*). Briefly, the mean within-module value for each brain map vector was calculated and compared to a null distribution (spin tests after excluding modules of fewer than ten nodes). The *z* values were transformed into *p* values and *p* < 0.05 applied to determine statistical significance. In the margins of *j*, we show the number of maps that were significantly enriched for each module.

**FIG. S13.**
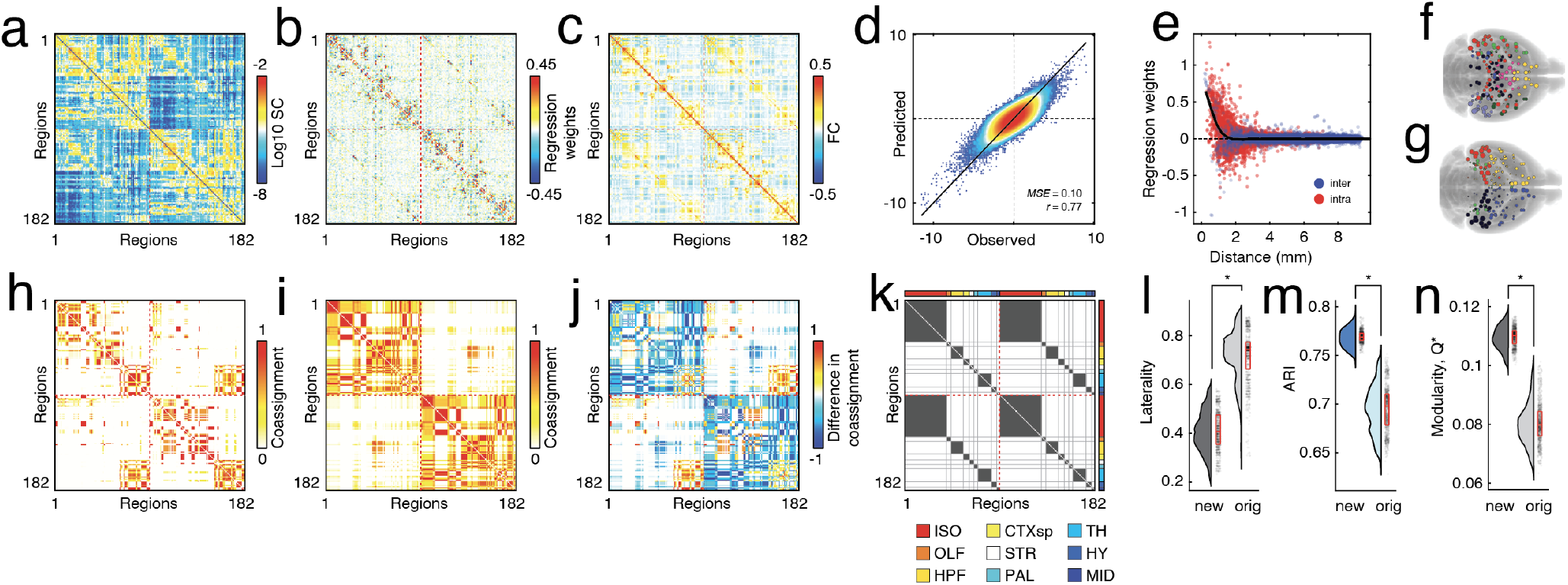
Replication of main effects using hyper-dense tract-tracing data from mice. (*a*) Logarithm of anatomical connection weights. (*b*) Estimated weights using linear model. (*c*) Functional connectivity (correlation). (*d*) Two-dimensional histogram of predicted and observed activity across all regions and mice. (*e*) Scatterplot of estimated edge weights versus Euclidean distance. (*f*) Communities estimated by applying modularity maximization to the estimated edge weight matrix. (*g*) Communities estimated by applying modularity maximization to the anatomical connectivity matrix. Panels *h* and *i* show module coassignment matrices for the two versions of edge weights. Panel *j* shows the difference in edge weights. (*k*) Areal coassignment matrix. Panels *l-n* show laterality of detected communities, similarity of detected partitions with respect to areal labels (adjusted Rand index), and the modularity of the detected partitions.

**FIG. S14.**
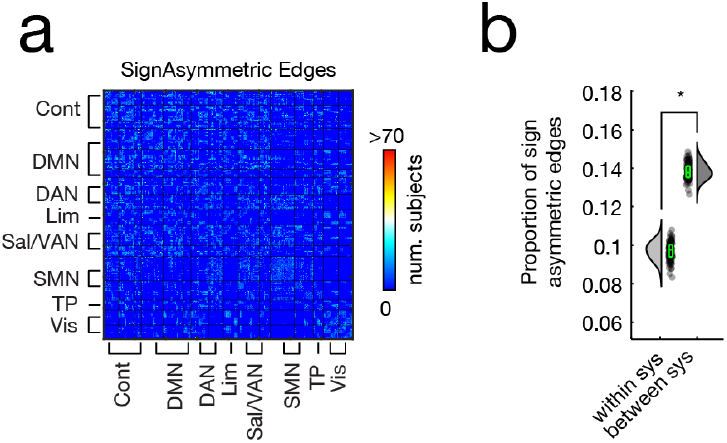
Signed asymmetry between incoming and outgoing edges is most common between functional brain systems. (*a*) The number of sign asymmetric edges across 95 subjects organized by system. (*b*) Proportion of existing edges that exhibit sign asymmetry within systems *versus* between systems.

**FIG. S15.**
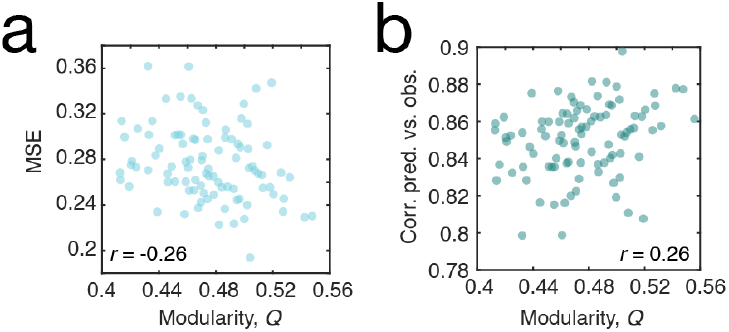
Relationship between fitness of the linear model and modularity of the functional connnectivity matrix. (*a*) Relationship between modularity (Q) and mean square error (MSE). (*b*) Relationship between modularity (Q) and the correlation between predicted and observed time series.

